# Continuous nuclear envelope surveillance is required for DNA double-strand break repair

**DOI:** 10.1101/2024.07.12.603222

**Authors:** Sara Medina-Suárez, Félix Machín

## Abstract

Precise double-strand break (DSB) repair is paramount for genome stability. Homologous recombination (HR) is preferred to repair DSBs when a nearby sister chromatid ensures an error-free template. In *Saccharomyces cerevisiae*, this preference extends into anaphase and telophase (late mitosis; late-M), despite sister chromatids having been pulled apart. Previously, we identified the nuclear envelope (NE) protein Msc1 as important for late-M DSB repair. Here, we report that Msc1 faces the NE lumen, and its depletion leads to DSB-independent over-compartmentalization of the nucleus, which hampers approximation of sister loci after DSB. Depletion of Msc1 also leads to nuclear pore complex mislocation, a phenotype shared by the highly conserved NE healing complex ESCRT-III. Critically, we show that both Msc1 and ESCRT-III also ensure DSB repair in G2/M, and that there is synergism between Msc1 and the ESCRT-III subunit Snf7. These findings highlight the essential role of NE health in DSB repair.

## Introduction

DNA double-strand breaks (DSBs) jeopardize cell survival and genome stability, crucially contributing to carcinogenesis ^1,2^. Cells employ two primary DNA repair mechanisms against DSBs: non-homologous end joining (NHEJ) and homologous recombination (HR). NHEJ relies on error-prone pathways to mend broken DNA ends, while HR utilizes intact homologous sequences for precise restoration. The choice between NHEJ and HR hinges on cyclin-dependent kinase (CDK) activity, dictating preference based on the cell cycle phase ^3–5^. HR demands a well-aligned sister chromatid to be error-free, which makes it optimal for S and G2 phases. However, CDK activity remains high well into mitosis (M phase), even after anaphase onset breaks down the proper alignment of sister chromatids. In complex eukaryotes, extensive chromosomal condensation in early M phase impedes HR ^6–8^. Conversely, in simpler eukaryotes like yeast, HR remains active in late mitosis (late-M; anaphase/telophase) and appears to repair some DSBs through re-alignment of segregated sister chromatids ^8–10^. One crucial step for this realignment is a change in the late-M dynamics of the microtubule cytoskeleton, both the spindle and astral microtubules, that forces the relaxation of the elongated nucleus and thus the re-approximation of the segregated sisters ^10^. This way, the ends of the DSB might realign with the unbroken sister for an effective HR-driven DSB repair.

An important yet frequently overlooked event during late-M is the massive remodeling of the nucleus. Eukaryotic chromosomes are surrounded by the nuclear envelope (NE), a double membrane that is continuous with the endoplasmic reticulum (ER). The NE can either remain intact or disassemble prior to chromosome segregation, defining two major forms of mitosis termed closed and open mitosis, respectively ^11^. Closed mitosis is considered the most ancient form and is found in many fungi and unicellular protists, including important human parasites. In contrast, most plant and animal cells undergo open mitosis. Although the NE remains during the closed mitosis, it still undergoes significant changes. In particular, its shape transitions from a sphere to an hourglass in early anaphase and to a dumbbell in late anaphase. In the latter morphology, a thin bridge of nucleoplasmic material connects the segregating nuclei until cytokinesis ^12^. Cytokinesis is preceded by karyokinesis, yet how this mechanistically occurs and is coordinated with the transition to a new cell cycle is poorly understood. In the fission yeast *Schizosaccharamyces pombe*, the NE protein Les1 has a direct role in modulating karyokinesis by ensuring the local corralling of a subset of nuclear pore complexes (NPCs) at the mid-bridge ^13^. This appears to correctly position karyokinesis when these NPCs disassemble in late-M to trigger a local NE breakdown reminiscent of the general NE breakdown observed in open mitosis ^13,14^. Completion of karyokinesis must then occur through membrane fission/fusion events, a process that is likely executed by the highly conserved ESCRT-III complex ^13,15^. This complex also deals with many other physiological and accidental wounds to the NE that pose a risk for cell survival or genetic stability as nucleoplasm leaks into the cytoplasm and vice versa ^15^. In fact, the knockout mutant for SpLes1 not only causes karyokinesis malposition but also leads to transient NE ruptures that must be repaired by ESCRT-III to preserve cell viability ^13,16^. Another role of ESCRT-III on the yeast NE homeostasis is the control of the quality and distribution of NPCs ^17^, which suggests not only a tight link with the nucleus-cytoplasm shuffling but also with DSB repair. Indeed, eroded telomeres and other one-ended DSBs are recruited to NPCs to be repaired by the specific HR subpathway known as break-induced replication (BIR) ^18,19^. NPCs also tether DSBs that are difficult to repair by HR and undergo NHEJ instead ^20^.

In a previous work, we found through proteomics that the only SpLes1 ortholog in *S. cerevisiae*, Msc1, plays a particular role in facilitating DSB repair in late-M ^21^. In the present work, we show that Msc1 controls proper NE shape and NPC distribution, which establishes a novel connection between NE homeostasis and DSB repair in late-M. In addition, we also show that the highly conserved NE healing complex ESCRT-III partly phenocopies these defects and cooperates with Msc1 in ensuring cell recovery from DSBs.

## Results

### Msc1 faces the NE lumen and does not translocate after DNA damage

Msc1 is the only putative ortholog in *S. cerevisiae* of two in *S. pombe* proteins, Les1 and Ish1. In general, little is known about the function of these three proteins; however, Ish1 and Les1 are known to be distributed along the inner and outer nuclear membranes (INM and ONM) of the NE and to form homo- and heterodimers ^22^. Both Ish1 and Les1 are characterized by the presence of several Ish1 motifs (pfam PF10281), whose molecular function(s) is unknown, a putative C2H2 zinc finger (ZnF) DNA-binding domain, and a unique transmembrane (TM) domain at the very N-terminus (Fig 1a) ^13,22^. The predicted number of Ish1 motifs differs between papers, from 2 to 5-6, but 4 is the most likely number based on recent in silico predictions and motif alignments ^22^, with the third Ish1 motif overlapping with the C2H2 ZnF. Like its putative *S. pombe* orthologs, Msc1 is predicted to contain four Ish1 motifs (https://www.genome.jp/tools/motif/; independent E-values <10^-10^) and a single N-terminal TM domain. However, unlike SpLes1 and SpIsh1, the putative C2H2 ZnF in Msc1 does not overlap with any Ish1 motif (Fig 1a).

**Figure 1.**
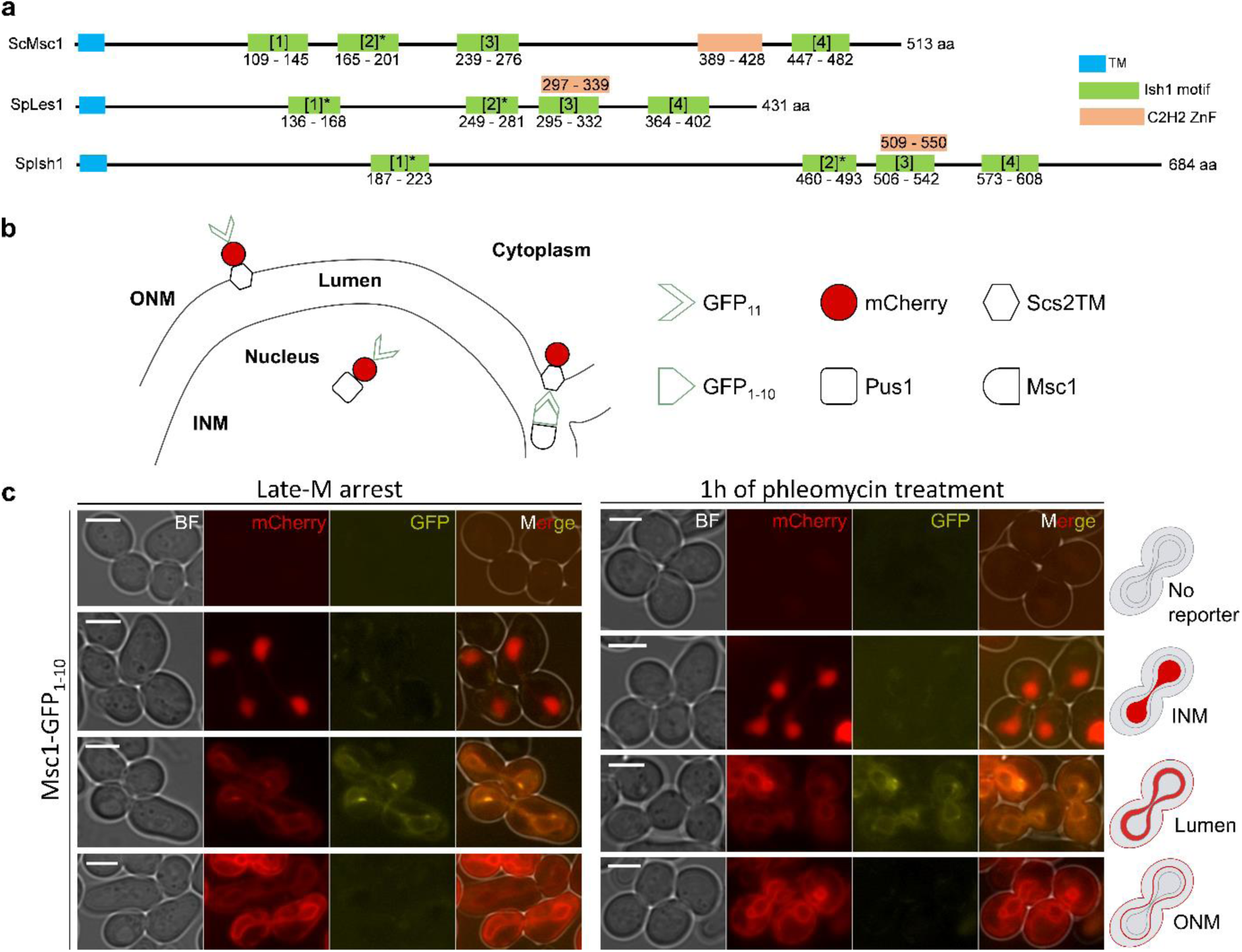
Spatial arrangement of Msc1 in the nuclear envelope. **(a)** Protein structure of ScMsc1 and its close orthologs SpIsh1 and SpLes1. The different protein motifs are depicted as boxes. Proteins and motif lengths are to scale, with delimiting amino acids numbered under or in the boxes. **(b)** Schematic of the system used to determine the NE face of the Msc1 C-terminus (modified from ^24^). The system is based on three strains with different mCherry-GFP_11_ chimeras reporting for INM facing the nucleoplasm (Pus1-mCherry-GFP_11_), ONM facing the cytosol (mCherry-GFP_11_-Scs2TM), and INM/ONM facing the NE lumen (Scs2TM-mCherry-GFP_11_). The GFP_11_ is a small portion (3 Kd) of the GFP that complements the larger GFP_1-10_ fragment (24 Kd), which in turn is tagged to the protein of interest (Msc1 in our case). Only if GFP11 binds to GFP1-10, a full and functional GFP is reconstituted. **(c)** Based on such a system, the spatial location of the functional globular tract of Msc1 was determined in the late-M NE both before and after DSB generation (1 h phleomycin). In all scenarios, reconstituted GFP was seen only in the strain that reports for the NE lumen. Representative micrographs are shown.

The TM domain determines that the N- and the C-termini of the protein must face different compartments. The close proximity of the TM to the N-terminus implies that all protein functions reside between the TM sequence and the C-terminus. Interestingly, the ultrastructural analysis of both SpIsh1 and SpLes1 by electron microscopy has shown that the C-terminal faces the lumen of the NE ^22^. This configuration has profound implications as, for instance, the putative C2H2 ZnF DNA-binding domain cannot interact with the chromatin. It is worth noting that C2H2 ZnF proteins can organize chromosome architecture ^23^, so Msc1 could directly modulate chromosome organization during DNA damage provided that its C2H2 ZnF faces the nucleoplasm. Likewise, the Ish1 motifs share similarity with the conserved HeH/LEM (pfam PF12949) and SAP (pfam PF02037) domains ^13^, which have DNA-binding and chromatin-anchoring properties as well.

To check the spatial configuration of Msc1, we used a system based on the reconstitution of a split GFP when a target protein and a localization reporter reside in the same compartment ^24^. We C-terminal tagged Msc1 with one half of the GFP (GFP_1-10_) and introduced the construct into three strains that differ in the localization of the other half of the GFP reporter (GFP_11_) (Fig 1b). One reporter, Pus1, is set for the nucleoplasm and allows GFP reconstitution when the protein of interest is located in the nucleoplasm or facing the nucleoplasm from its location at the INM. A second reporter, Scs2, is an integral ONM/ER protein with the N-terminus facing the cytoplasmic side and the C-terminus facing the space between INM and ONM (lumen). In this manner, N- vs C-terminal tagging with GFP_11_ makes Scs2 a reporter for ONM/cytoplasm and lumen, respectively. In addition to the GFP_11_ half, all these reporters carry a fused mCherry as an internal control (Fig 1b). To synchronize the cell culture in late-M, we used the thermosensitive *cdc15-2* allele of the kinase Cdc15, which is critical for the telophase-G1 transition. At 34 °C, this mutant arrests cells in late-M with sister chromatids segregated and the NE elongated in hourglass/dumbbell shapes ^12,25,26^. Of the three reporters, GFP reconstitution with Msc1-GFP_1-10_ was only possible with the Scs2TM where the GFP_11_ was facing the NE/ER lumen (Fig 1c, left panels). This implies that the putative C2H2 ZnF domain cannot be in contact with the chromatin during the late-M arrest. However, it could still be possible that Msc1 translocates or flips exclusively upon the generation of DSBs, resulting in the functional part of the protein, including the ZnF, facing the nucleoplasm. There are several ways of generating DSBs. A simple and practical way is treating the cells with phleomycin, a drug that causes DSBs at random location through a mechanism that mimics that of ionizing radiation. Thus, we performed GFP reconstitution experiments after DSB generation by phleomycin, yet we still found that Msc1 was entirely facing the lumen (Fig 1c, right panels).

### Loss of Msc1 results in aberrant NE morphologies in late mitosis

The fact that Msc1 always faces the NE lumen implies that its role on DSB repair must be indirect; i.e., Msc1 physically interacts with neither DSBs nor HR factors. SpLes1 and SpIsh1 have been proposed as a modulator of karyokinesis ^13,14^. When SpLes1 is depleted, karyokinesis is misplaced and the NE is transiently damaged ^13^. In this context, we reasoned that the nucleoplasmic bridge of *S. cerevisiae* cells could become compromised in Δ*msc1* during the protracted arrest in late-M that precedes DSB generation. For instance, premature karyokinesis in *cdc15-2* Δ*msc1* could irreversibly preclude the search for the sister chromatid during the partial anaphase regression. Hence, we sought signs of premature karyokinesis in Δ*msc1*. For this purpose, we labeled the NE with Sec61 and Nup49. The former is an NE/ER protein that continuously and more strongly labels the NE, whereas the latter is an NPC component that specifically labels the NE, although it tends to give a more punctate (and thus imprecise) pattern ^27^. In general, signs of premature karyokinesis in Δ*msc1*, seen as a partial or complete absence of the bridge, were rather modest or absent. A Sec61 bridge was observed in ∼95% of WT cells and ∼90% of Δ*msc1* cells, whereas bridging Nup49 was observed in similar proportions in both strains (∼80%) (Fig 2a and S1a; mock experiments). In addition, we also directly examined the continuity of the nucleoplasmic bridge using the freely circulating nucleoplasmic TetR-YFP reporter. With this marker, we found only a modest increase (∼10%) of signs that can be compatible with premature karyokinesis in Δ*msc1* cells (Fig S1b). Next, we looked at whether karyokinesis was instead accelerated during DSB generation, as the stress caused by DNA damage and the general mobilization of the chromatin for its repair may destabilize the NE bridge in Δ*msc1*. In this case, and in addition to phleomycin, we also used a pair of DSBs generated at a single location on chromosome III, one per sister chromatid. The system to generate these DSBs is based on the controlled expression of the HO endonuclease, which cuts the *HOcs* sequence at the *MAT* locus ^28,29^. With either strategy, a similarly high percentage of cells retain the bridge (Figs 2a and S1a,b).

**Figure 2.**
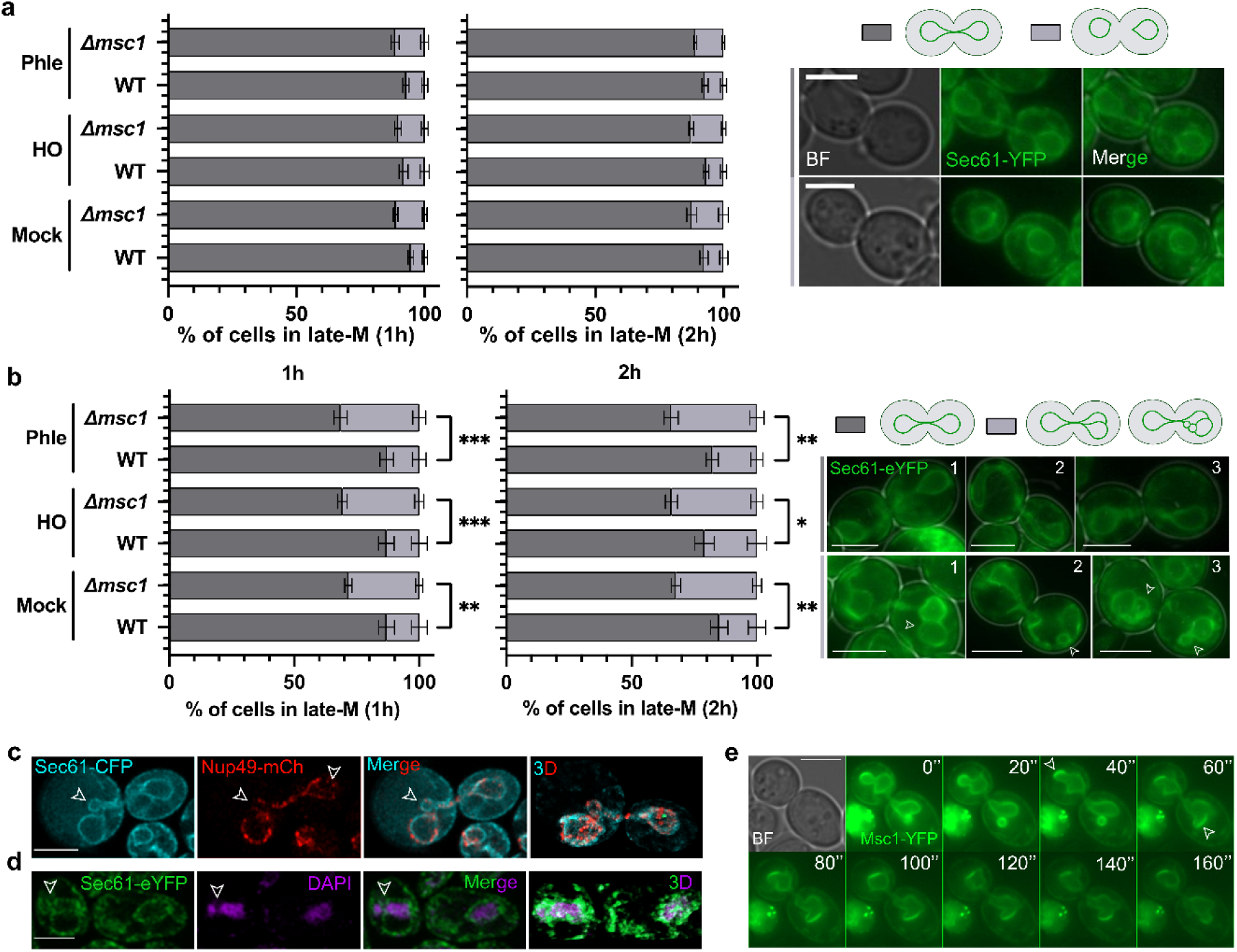
Role of Msc1 in late-M NE shape. (**a**) Karyokinesis status as reported by the presence (dark gray bars) or absence (light gray bars) of a Sec61-eYFP NE bridge in WT and *Δmsc1* late-M cells in the absence (mock) and presence of DSBs generated by either phleomycin (phle) or HO (mean ± s.e.m., n=3). Representative micrographs of the mock experiment are shown on the right. **(b)** Abnormal NE shapes, including blebs and partitions, observed in the same experiment (mean ± s.e.m., n=3). On the right, three representative micrographs of normal dumbbell late-M shapes (top), and three representative micrographs of abnormal shapes (bottom) with partitions indicated by arrowheads. **(c,d)** High resolution of nuclear partitions in *Δmsc1* late-M cells. In segmented nuclei, the NE septum contains NPC and nuclear partitions contain DNA. **(c)** Cells expressing Sec61-eCFP and Nup49-mCherry were fixed and visualized by confocal microscopy with airyscan 2 superresolution. A central slice of a representative cell with one of the segregating nuclear bodies presenting a partition indicated by the arrowhead (left) and a 3D reconstruction of the same cell (right). **(d)** Cells expressing Sec61-eYFP were stained with DAPI and visualized as above. A central slice of a representative cell with a partition indicated by the arrowhead that contains DNA in both lobes (left). 3D reconstruction of the same cell (right) **(e)** Msc1 forms patches at NE blebs. A short time-lapse series of Msc1-eYFP after the DSB generation. Note how Msc1 concentrates on NE blebs (arrowheads) and dynamically corrects this aberration. Scale bars correspond to 3 μm; BF, bright field. The unpaired t test was used for statistical comparisons (*** for p<0.001, ** for p<0.01 and * for p<0.05).

Even though premature karyokinesis was not observed in Δ*msc1*, during the course of these experiments we notice that a higher proportion of cells presented aberrant morphologies in one or both segregated nuclear bodies. These aberrations included nuclear bulges and what appeared to be partitions within the nucleus. Although some of these aberrant shapes were already observed in *cdc15-2* arrested cells, the percentage in late-M was significantly higher in Δ*msc1* (Fig 2b; ∼15% in WT vs ∼30% in Δ*msc1*). Airyscan superresolution analysis of Δ*msc1* co-expressing Sec61-CFP and Nup49-mCherry revealed that the septum separating the apparently partitioned nucleus is in fact made of NE (∼90% of these cells have a septum where Sec61 and Nup49 colocalize, n=237 cells) (Fig 2c, S2 and movie S1). Strikingly, the septum often appeared to be closed, as far as the airyscan superresolution can determine. More strikingly, both compartments contain DNA in nearly 90% of these examples (n=143 cells) (Fig 2d and S3). The proportion of aberrant late-M nuclei did not change after DSB generation in *MSC1* and Δ*msc1* strains (Fig 2b).

In the previous work, we reported that Msc1 is not uniformly localized in the NE but forms concentrated patches, and there is also a correlation between the presence of patches and DSB repair factories ^21^. When we looked at abnormal nuclear shapes in the *MSC1* strain after DSB generation, we found that Msc1-YFP appears enriched in NE bulges, and short time lapse movies suggest that Msc1 may participate in correcting these bulges, thus re-establishing a rounder NE and preventing septation (Fig 2e and S4).

### Msc1 favors back migration of sister chromatids during DNA repair in late mitosis

One logical consequence of the observed nuclear aberrations is that a proportion of late-M cells, which is higher in Δ*msc1*, is physically disabled from finding the correct template for HR once DSBs occur. Since this step requires the back migration of sister loci through the bud neck ^10^, we compared both back migration and coalescence events in Δ*msc1* versus WT after phleomycin (Fig 3). In the WT, we observed a significant increase in these events at the right telomere of chromosome XII (cXIIr-Tel; from 5% to 15% in late-M cells after phleomycin). However, this increase was about half in Δ*msc1* (Fig 3a-c). Likewise, the formation of retrograde chromatin bridges after phleomycin was largely impaired in Δ*msc1* (Fig 3d-f). Chromatin was visualized through the histone A2 (Hta2-mCherry), and it went from more than a 2-fold increase in bridges in the WT after DSB generation (from ∼25% to ∼60%) to half that increase in Δ*msc1* (from ∼20% to 30%). These experiments confirmed that Δ*msc1* late-M cells are less capable of bringing together the segregated sister chromatids upon DSBs.

**Figure 3.**
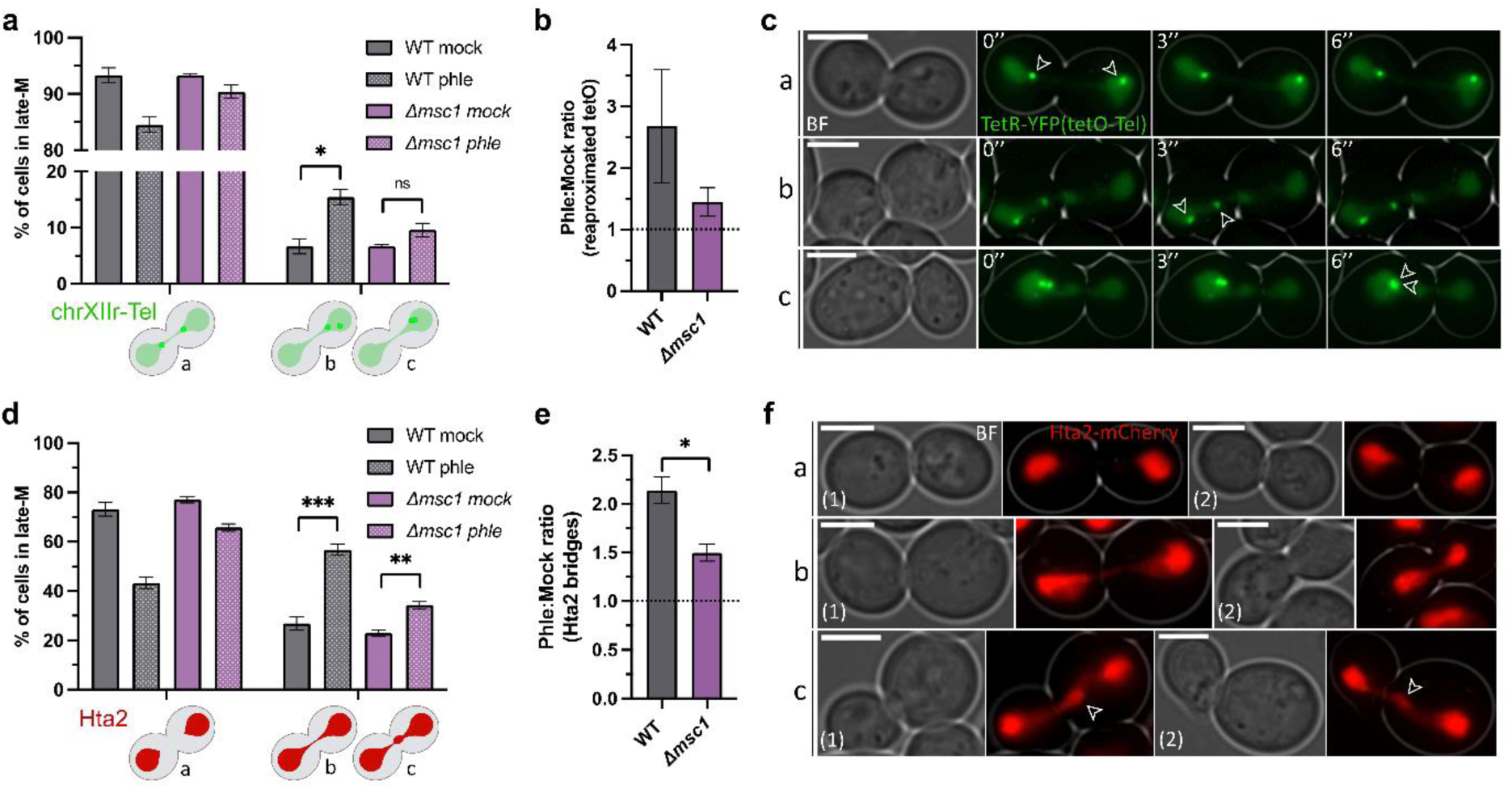
Back migration and coalescence of sister loci is hampered in *Δmsc1*. **(a)** WT and *Δmsc1* cells carrying a TetR-YFP/*tetOs* construct that labels the right telomere of chromosome XII (cXIIr-Tel) were treated as in Figure 2 (phleomycin only). Samples were taken 1 h after phle (or mock) addition, and late-M cells classified in three categories: a, segregated sister cXIIr-Tels; b, back migration of one sister cXIIr-Tel towards the other; c, coalescence between sister cXIIr-Tels (mean ± s.e.m., n=3). Categories b and c are represented together because they are rapidly interchangeable. **(b)** Phle:mock ratios of cXIIr-Tel retrograde events in WT and *Δmsc1* late-M cells. (**c**) Representative micrographs of the three quantified categories. A short frame of a 6’’ movie is shown to better appreciate the dynamism of coalescence (bottom line). Arrowheads point to the cXIIr-Tel sisters. (**d**) Like in panel (a) but with strains that label the bulk of chromatin with Hta2-mCherry (mean ± s.e.m., n=3). Late-M cells were classified in three categories again: a, no chromatin bridge; b, thin bridge; c, bridge with bulges. Categories b and c are represented together. **(e)** Phle:mock ratios of retrograde chromatin bridges in WT and *Δmsc1* late-M cells. (**f**) Representative micrographs of the three quantified categories. Two examples of each are shown. Arrowheads point to a chromatin bulge often seen in retrograde bridges. Scale bars correspond to 3 μm; BF, bright field. The unpaired t test was used for statistical comparisons (*** for p<0.001, ** for p<0.01, * for p<0.05, and n.s. for p>0.05).

We have previously shown that DSBs in late-M exert a partial regression of anaphase, with shortened distances between segregated sister centromeres, dismantlement of the anaphase spindle and redistribution of the spindle motor type 5 kinesin Cin8 towards the poles^10^. Thus, we also checked whether these anaphase regression phenotypes were affected in Δ*msc1*. We looked at these three markers of regression, observing no differences between the *MSC1* and the Δ*msc1* strains before and after DSB generation (Fig S5). First, the anaphase spindle (as reported by GFP-Tub1) was seen in 70-80% of late-M cells, whether *MSC1* or Δ*msc1* (Fig S5a). DSB generation by phleomycin caused a drop in these percentages to 30%, though it occurred both in the WT and Δ*msc1*. A similar pattern was seen for Cin8-mCherry; nearly 90% was found on the spindle in late-M cells for *MSC1* or Δ*msc1* cells, with an abrupt drop to just 20% after phleomycin, with no distinction between *MSC1* or Δ*msc1* (Fig S5b). Lastly, the distance between sister chromosome XII centromeres was shortened after phleomycin, but to a similar extent in *MSC1* or Δ*msc1* (Fig S5c). Thus, we concluded that the reduction in chromatin bridges and coalescence of sister loci at chromosome arms is not due to the inability to relieve the late anaphase segregating tension by the spindle apparatus. This conclusion supports in turn the potential role of NE septation in this defect.

### The NE maintenance complex ESCRT-III is important for DSB repair

In addition to its role in karyokinesis, SpLes1 and SpIsh1 might have a more general role in the NE homeostasis. For instance, the nuclear-cytoplasmic barrier is compromised in *les1*Δ, which is in turn synthetically lethal with several components of the evolutionarily conserved ESCRT-III complex in *S. pombe* ^13,16^. The basis of this extreme negative genetic interaction is that ESCRT-III repairs NE wounds that occur in the absence of SpLes1 ^13^. Indeed, this is one of the most important functions of ESCRT-III in eukaryotes, i.e., surveys NE integrity and seals unwanted NE pores ^15,30^. We envisioned that a similar scenario might be taking place in *S. cerevisiae* without Msc1, with a likely aggravated circumstance due to the overstretched NE at late-M.

In *S. cerevisiae*, ESCRT-III comprises up to 8 subunits, with Snf7 being the core structural protein, and the Did4/Vps24 dimer forming a major regulatory player ^15,31^. The ESCRT-III machinery is completely absent in the Δ*snf7* mutant, whereas it is malformed and non-functional in either Δ*did4* or Δ*vps24*. To check whether ESCRT-III is involved in DSB repair, we determined sensitivity to phleomycin through both spot assays and growth curves (Fig 4a and S6). In all cases, loss of function of the complex led to hypersensitivity to phleomycin, with Δ*snf7* giving the strongest phenotype. In spot assays, Δ*snf7* phenocopied the sensitivity profile of Δ*msc1* in our YPH499 genetic background, which in turn phenocopied that of Δ*rad52* (Fig 4a and S6). Interestingly, in the BY4741/S288C background, phleomycin hypersensitivity of Δ*snf7* was even stronger (Fig S7). The other two ESCRT-III mutants gave a milder, yet consistent, sensitivity in both genetic backgrounds.

**Figure 4.**
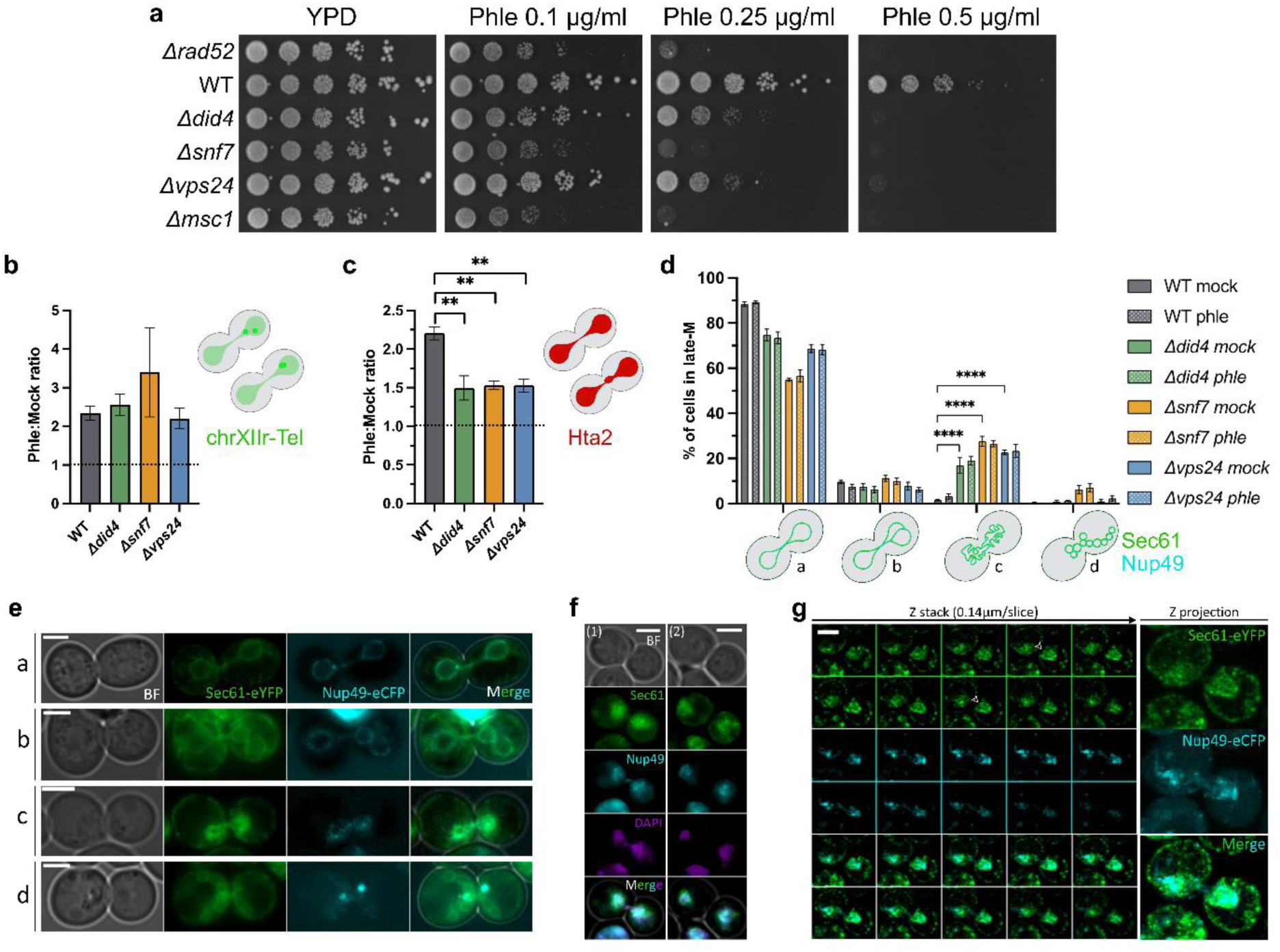
Mutants for the NE healing complex ESCRT-III partly phenocopy *Δmsc1*. **(a)** Spot assay against phleomycin of ESCRT-III mutants. The HR mutant *Δrad52* and *Δmsc1* were included as references. Note that the *Δsnf7* (the core component of ESCRT-III) is as sensitive to DSBs as *Δrad52* and *Δmsc1*. (**b**) Phle:mock ratios of cXIIr-Tel retrograde events in WT and ESCRT-III-deficient late-M cells (mean ± s.e.m., n=3). (**c**) Phle:mock ratios of retrograde chromatin bridges in WT and ESCRT-III-deficient late-M cells (mean ± s.e.m., n=3). (**d**) Abnormal late-M NE morphologies in ESCRT-III mutants (mean ± s.e.m., n=3). Four categories were assessed based on Sec61-eYFP: a, normal hourglass shape; b, partitioned; c, blurred; d, multivesiculated. The strain also carries the Nup49-eCFP construct. (**e**) Representative micrographs of the four categories. (**f**) Representative micrographs of the nuclear DNA staining (DAPI) pattern in a blurred late-M NE. Two examples are shown; (1) with close nuclear masses, and (2) with more distant masses. (**g**) Confocal images with airyscan superresolution of a representative blurred late-M NE. On the left, the sequence of z planes; on the right, the 2D projection. The arrowhead points to a possible septum. Scale bars correspond to 3 μm; BF, bright field. The unpaired t test was used for statistical comparisons (**** for p<0.0001, ** for p<0.01).

### ESCRT-III preserves the NE in late mitosis and facilitates anaphase retrograde events after DSBs

To check whether the DSB hypersensitivity of ESCRT-III mutants was mechanistically related to that of Δ*msc1*, we next phenotypically examined late-M cells after DSBs in these mutants. Firstly, we quantified retrograde events for cXIIr-Tel, and found that the phleomycin vs. mock ratio was similar between the WT and the mutants (∼2.5); and it might be even higher in Δ*snf7* (Fig 4b and S8a). On the contrary, the increase in retrograde chromatin bridges (Hta2-cCherry) after phleomycin was more modest in the ESCRT-III mutants relative to the WT (Fig 4c and S8b). Thus, with these mutants, we found for the first time a loss in the positive correlation between sister telomere retrograde events and the presence of de novo chromatin bridges. One possible explanation for this is that only the retrograde movement of chromatin not bound to the NE is affected since cXIIr-Tel is attached to the NE. However, it is important to note that this is based on the comparison of phle:mock ratios. In terms of global cXIIr-Tel retrograde events after phleomycin, there were fewer in in Δ*snf7* as well (Fig S8a).

Next, we checked NE morphologies in these mutants to determine if they phenocopied that of Δ*msc1*. Unexpectedly, inner septation was as rare as in the WT strain (Fig 4d,e). Instead, two other aberrant morphologies were observed. On the one hand, around ∼20-30% of late-M cells presented blurred Sec61 and Nup49 signals, which gave these extended nuclei the appearance of being formed by discontinuous patches of NE (Fig 4e, category “c”). On the other hand, in ∼5% of late-M Δ*snf7* cells, Sec61 appeared multivesiculated (Fig 4e, category “d”). These aberrant morphologies preceded the induction of DSBs and their relative percentages remained after adding phleomycin (Fig 4d). Thus, if these NE aberrations affect DSB sensitivity, they likely yield an inadequate nuclear architecture for later DSB repair. Importantly, and despite Sec61 and Nup49 could not outline the NE, their signal perfectly colocalized with the nuclear DNA, which appeared normal under the microscope (Fig 4f and S9). To better determine the structural feature of this abnormal NE, we drew again upon airyscan superresolution microscopy (Fig 4g and S10). However, we could little improve the resolution of these aberrations, with Sec61 appearing spread over the nucleus in confocality. One explanation for this pattern is that the NE contains multiple invaginations, which would also fit with what has been reported in human cells ^32^. Airyscan superresolution also outlined signs of inner septation (Fig 4g and S10, arrowheads), albeit they were less defined than in Δ*msc1*.

### ESCRT-III core component Snf7 is synergistic with Msc1 in DSB repair and late-M DSB-associated events

Having shown that Δ*msc1* and ESCRT-III mutants, especially Δ*snf7*, individually hypersensitized to DSBs, and that they both coincide in having aberrant yet distinct late-M nuclei, we next tested whether they together synergically change these phenotypes. Noticeably, *S. pombe les1* is synthetically lethal with deletions in ESCRT-III genes ^16^. Thus, to combine depletion of Msc1 and Snf7 in the same strain we opted for tagging Snf7 with the auxin-inducible degron minimal sequence (AID*) in the Δ*msc1* strain. Snf7-AID* was quickly degraded after adding the auxin indol-acetic acid (IAA), and no protein could be detected by Western blotting after 1 hour (Fig 5a). We then tested whether there exists a negative genetic interaction between *MSC1* and *SNF7*. In spot assays, we found that the strain carrying Δ*msc1 SNF7-AID* OsTIR1* (OsTIR1 is required for auxin to target AID* for degradation ^33^) was unaffected by IAA, and cells grew as well as in isogenic strains without the OsTIR1 (Fig 5b). This result is thus in marked contrast with what has been reported in *S. pombe* ^16^. Likewise, a complete lack of negative genetic interaction was seen for Δ*msc1* and depletion of either Did4 or Vps24 (Fig S11a,b).

**Figure 5.**
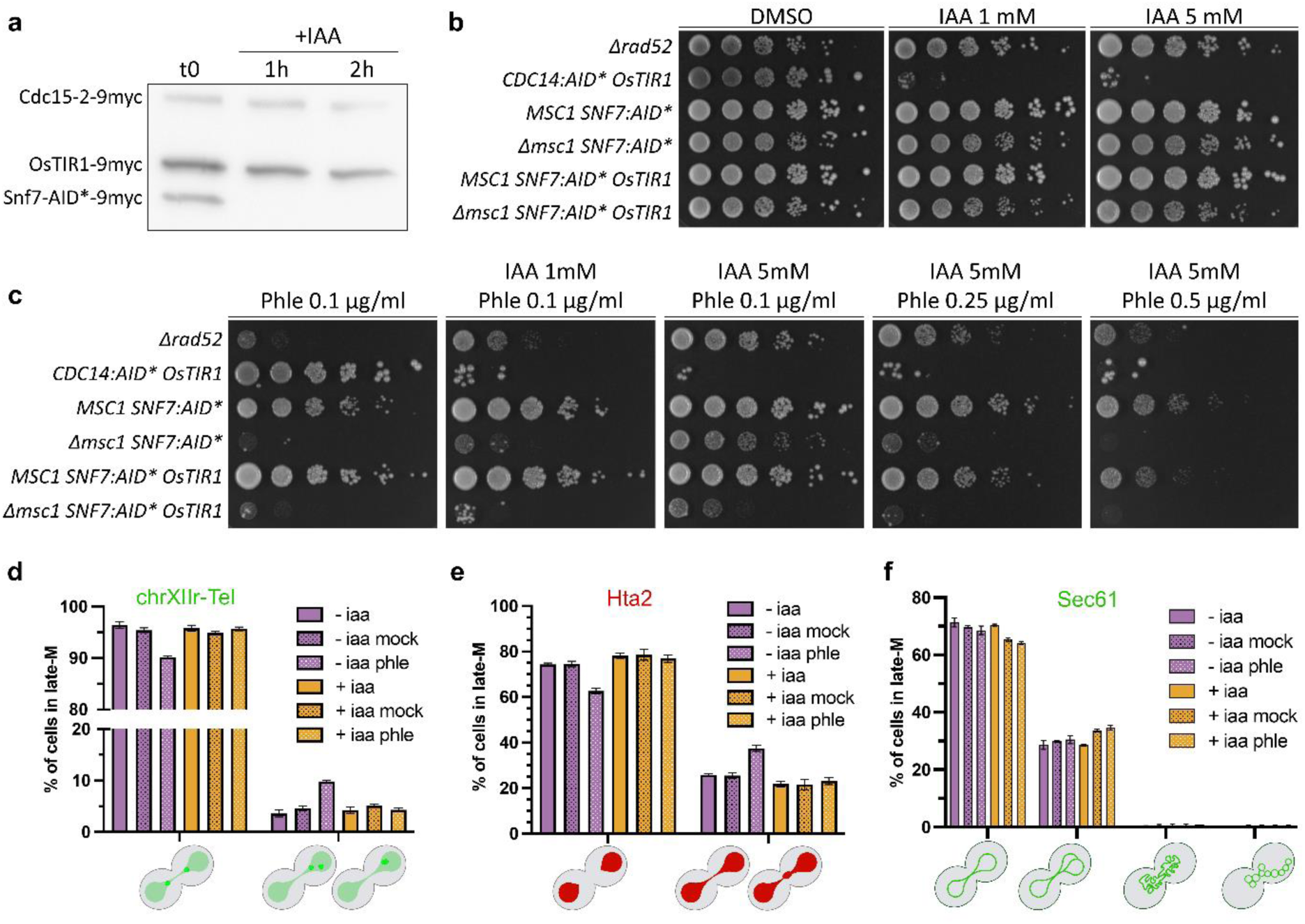
Synergism profiles of Msc1 and ESCRT-III in DSB sensitivity and late-M events. **(a)** Western blot to show that the core ESCRT-III component Snf7 tagged with the AID* degron is degraded upon auxin addition (+IAA). The blot was probed with an anti-myc antibody, which recognizes three tagged proteins in this strain: Snf7-AID*-9myc proper, Cdc15-2-9myc and OsTIR1-9myc. The latter two served as loading controls. **(b)** Spot assay against IAA of ESCRT-III *Δmsc1* double mutants. The *CDC14:AID* OsTIR1* strain was included as a control for IAA. *MSC1* and just *SNF7:AID\** (no OsTIR1) strains were also included as a reference for the putative genetic interaction and to assess the effect of Snf7 C-terminal tagging, respectively. **(c)** Spot assay against phleomycin of ESCRT-III *Δmsc1* double mutants. The *Δrad52* strain was included as a control for phleomycin. *MSC1* and just *SNF7:AID\** (no OsTIR1) strains were also included as a reference for the putative genetic interaction and to assess the effect of Snf7 C-terminal tagging, respectively. **(d)** The segregation status of cXIIr-Tel in the *Δmsc1 SNF7-AID* OsTIR1* strain with and without depleting Snf7 (+/- iaa) and with and without DSBs (phle vs. mock) (mean ± s.e.m., n=3). Categories as in Figure 3a. The solid color bars (- iaa and + iaa only) represent the status at the late-M arrest, before splitting the culture into two for the mock and phleomycin treatments. (**e**) Like in panel (d) but following the chromatin with Hta2-mCherry (mean ± s.e.m., n=3). Categories as in Figure 3d. (**f**) Like in (d) but following abnormal late-M NE morphologies through Sec61-eYFP and Nup49-eCFP (mean ± s.e.m., n=3). Categories as in Figure 4d.

Next, we investigated through spot assays if a negative effect in these double mutants occurred upon DNA damage with phleomycin (Fig 5c, S11 and S12). We found that, in conditions where DNA damage did not increase the sensitivity of *MSC1* cells depleted of Snf7-aid* relative to cells having both Msc1 and Snf7 (*MSC1 SNF7-AID* OsTIR1* vs. *MSC1 SNF7-AID\**; 5 mM IAA, Phle 0.05-0.1 μg/mL), cells defective in both proteins (Δ*msc1 SNF7-AID* OsTIR1*) were more sensitive than those only defective in Msc1 (Δ*msc1 SNF7-AID\**) (Fig 5c and S12). Thus, these spot assays point out that there is a synergistic effect of depleting both proteins in relation to sensitivity to DSBs. However, this genetic interaction was not observed when depleting the two other ESCRT-III subunits, Did4 and Vps24 (Fig S11c,d) Incidentally, the combination of auxin and phleomycin diminishes the relative toxicity of the latter in all strains (Fig 5c, S11 and S12).

To check whether synergistic effects are also present for retrograde events in late-M, we determined approximation of sister cXIIr-Tel loci and retrograde chromatin bridges after phleomycin, and compared Δ*msc1* cells depleted or not of Snf7-aid* (Δ*msc1 SNF7-AID* OsTIR1* strain with or without IAA) (Fig 5d,e). For these experiments, the late-M cultures were first split in two and one of the subcultures treated with IAA for 1h to degrade Snf7-aid* (-iaa or +iaa), taking a sample as a reference for basal levels of retrograde events. Then, each culture was split in two again to add phleomycin to one of the new subcultures. All four subcultures (mock vs. phle) were then incubated as in other previous experiments. In this experimental condition, the reference culture just without Msc1 (-iaa subcultures) had a behavior similar to Δ*msc1* strains in previous experiments, with a ∼2- and ∼1.5-fold increase in cXIIr-Tel approximation and chromatin bridges, respectively, in phleomycin relative to the mock (Fig 5d,e). However, all retrograde events were fully prevented when cells were further depleted from Snf7 (+iaa subcultures). Thus, Msc1 and Snf7 (ESCRT-III) co-operate in bringing together sister loci in late-M after DSBs.

Lastly, we investigated whether there was also synergism in the aberrant NE structures seen for each mutant separately (Fig 5f). In this case, however, the septation phenotype observed in Δ*msc1* fully dominates over those seen after Snf7 depletion. Indeed, septation levels did not change after depleting Snf7 and, more interestingly, blurred Sec61/Nup49 were not seen after IAA addition. This latter result suggests that this phenotype might depend on Msc1 in cells depleted from Snf7. Alternatively, Snf7 depletion could have less penetrance than Δ*snf7* or, else, the blurred nucleus is the consequence of the cumulative absence of ESCRT-III through multiple generations, which cannot thus be unmasked by the short-term absence of Snf7 in this experiment.

### Late-M nuclear pore complex distribution is affected by Msc1 and ESCRT-III

Alongside the defects seen in the NE by Sec61, NPCs (Nup49) were largely mis-distributed and tend to concentrate in spots in the subset of cells affected by the blurred NE in ESCRT-III (Fig 4e-g and S9, S10). Interestingly, this uneven distribution was also observed in a proportion of cells in which the NE could be outlined (Fig 6a), so that the total percentage of late-M cells with an uneven distribution of Nup49 was as high as 60% in Δ*snf7* (and ∼40% in Δ*did4* and Δ*vps24*) compared to less than 10% in the WT *cdc15-2* (Fig 6b). The percentage of these aberrant nuclei did not change after generating DSBs with phleomycin.

**Figure 6.**
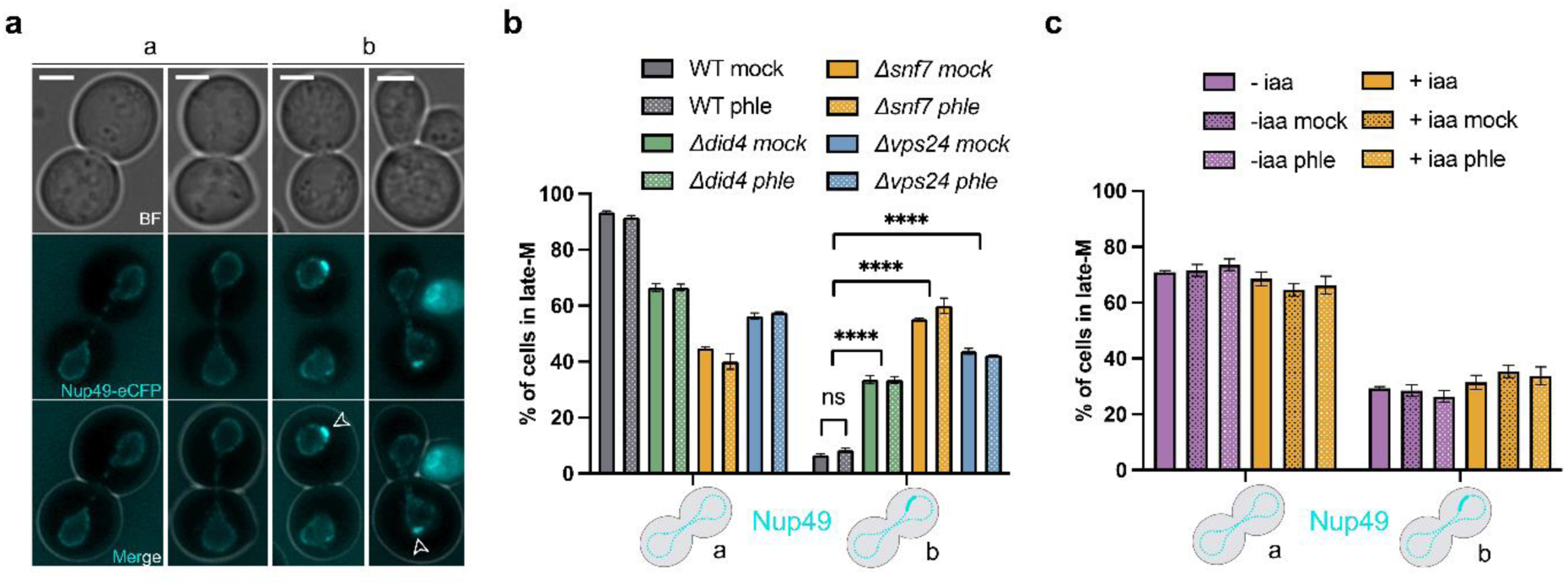
Nuclear pore complex distribution in strains depleted for the ESCRT-III complex and Msc1. **(a)** Representative micrographs of even (a) an uneven (b) distribution of Nup49 across the late-M NE. The arrowhead points to the characteristic patch of enriched Nup49 seen when distribution was uneven. Scale bars correspond to 3 μm; BF, bright field. **(b)** Late-M distribution of Nup49 in the WT and three ESCRT-III mutants with (phle) or without (mock) concomitant DSBs (mean ± s.e.m., n=3). **(c)** Late-M distribution of Nup49 in the *Δmsc1 SNF7-AID* OsTIR1* strain with and without depleting Snf7 (+/- iaa) and with and without DSBs (phle vs. mock) (mean ± s.e.m., n=3). The solid color bars (- iaa and + iaa only) represent the status at the late-M arrest, before splitting the culture into two for the mock and phle treatments.

To check whether this sick distribution of NPCs could be synergistically increased by the Msc1 depletion, we next looked at this phenotype in the Δ*msc1 SNF7-AID* OsTIR1* strain (Fig 6c). Strikingly, the Δ*msc1* mutation already rendered 20% of the late-M cells with an abnormal distribution, which did not change after generating DSBs. Similar to the results obtained for the aberrant blurred NE, the short-term depletion of Snf7 in this Δ*msc1* background neither increased Nup49 misdistribution nor led to the levels seen in Δ*snf7*.

### NE septation and NPC misdistribution also occur in G2/M-arrested cells in mutants for *MSC1* and the ESCRT-III complex

So far, we have studied phenotypes present in cells arrested in late-M since Msc1 was identified in a screening for DSB repair in this cell cycle stage ^21^. In asynchronous cells, DSBs activate the DNA damage checkpoint, which arrests cells in G2/M (metaphase-like) ^34^. Hence, we wondered what happened with the NE and NPCs during the phleomycin-driven G2/M arrest (Fig 7). We first checked whether the G2/M arrest was as effective in the mutants as is in the WT, observing similar cell percentages after 3h and 5h of phleomycin treatment (70-80%) (Fig 7a). In this G2/M arrest, cells appear mononucleated with either a round nucleus close to the bud neck or traversing it. In the latter case, the NE appears as bilobed with the neck making the constriction (bow-tie phenotype) (Fig 7b). When G2/M cells were released from DNA damage, there was a drop in viability for all mutants relative to the WT (Fig 7c). Only 60% of Δ*msc1* cells survived compared to the WT, whereas this percentage dropped to 20% for Δ*snf7*. Intermediate survival was obtained with the other two ESCRT-III subunits. This sensitivity profile fitted well with what was observed in spot assays above and emphasizes the importance that both Msc1 and ESCRT-III has in an effective DSB repair, beyond the particular cell cycle stage the cells are in when they receive the insult.

**Figure 7.**
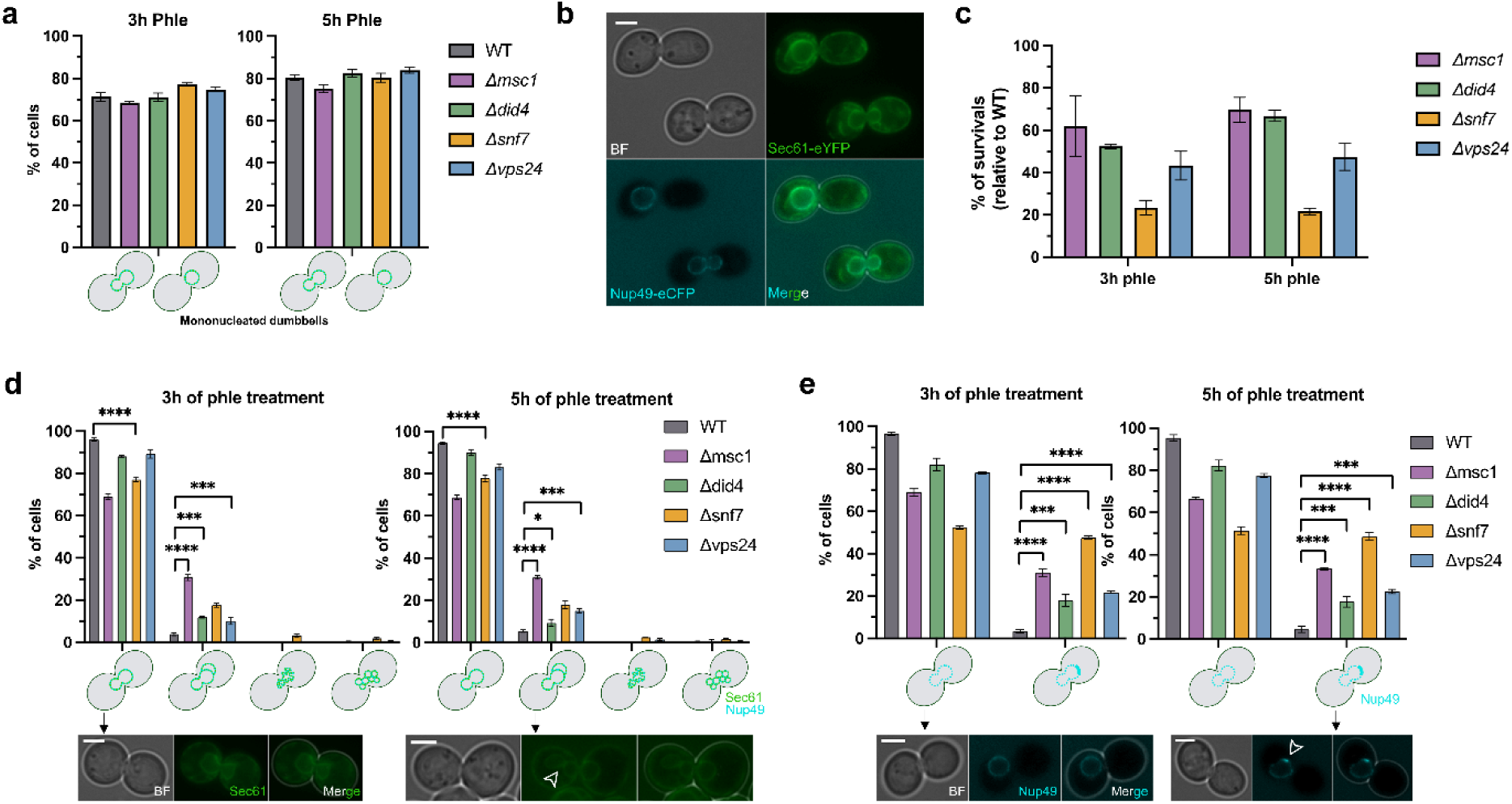
Nuclear envelope abnormalities are also present in a G2/M arrest elicited by DSBs. **(a)** Percentage of G2/M-arrested cells in the WT and ESCRT-III mutants after 3h and 5h of incubation with phleomycin. (mean ± s.e.m., n=3). Note that the ESCRT-III does not affect the arrest efficiency. G2/M-arrested cells were assessed by the presence of mononucleated dumbbell-shaped cells. The mononucleated nucleus could be either in one cell body or across the neck (bow-tie phenotype). **(b)** Representative micrograph with two G2/M- arrested cells with the two distinct NE shapes, round nucleus in one body (top) and with the bow-tie shape (bottom). **(c)** Cell viability of *Δmsc1* and ESCRT-III mutants after short-term G2/M blocks by DSBs (mean ± s.e.m., n=3). Percentage of survivors is relative to the isogenic WT. (**d**) Abnormal G2/M NE morphologies in *Δmsc1* and ESCRT-III mutants at the G2/M arrest elicited by DSBs (mean ± s.e.m., n=3). Four categories were assessed based on Sec61-eYFP; from left to right in the X-axis: normal round or bow-tie shapes; partitioned; blurred; multivesiculated. Representative micrographs for the normal and partitioned shapes are included underneath. The arrowhead points to the constriction that makes the partition. **(e)** Like in (d) but looking at the NPC (Nup49) distribution (mean ± s.e.m., n=3). Representative micrographs for the even and uneven distribution are shown below. Scale bars correspond to 3 μm; BF, bright field. ANOVA followed by the Dunnett’s test against WT was used for statistical comparisons (**** for p<0.0001, *** for p<0.001, * for p<0.05).

Next, we addressed the aberrant NE phenotypes described above in late-M. Septation was once again the major abnormality observed in Δ*msc1*, reaching up to 30% of G2/M cells (Fig 7d). Interestingly the ratio between the WT and the mutant was even higher than in late-M (∼7:1 after 3h in phle). Septation was also observed in the ESCRT-III mutants, especially in Δ*snf7*, although to a lesser extent. Interestingly, the other two phenotypes described in late-M, blurred and multivesiculated Sec61, were completely absent in G2/M (Fig 7d). Because these G2/M arrest experiments were performed at 25°C to keep Cdc15-2 active, whereas the late-M experiments were performed at 34°C, we checked the influence of the temperature in the NE in the Δ*snf7* mutant. We did not observe the blurred NE either (Fig S13). This points out that the blurred NE is a late-M specific phenotype of the Δ*snf7* mutant. On the other hand, G2/M NPC misdistribution was also observed for all mutants (Fig 7e). In this case, Δ*snf7* surpasses Δ*msc1* (∼50% vs. ∼35%), with values for both highly above the WT (<10%) and the other two ESCRT-III mutants (∼20%).

Finally, we investigated the subcellular location and abundance of Snf7 before and after DSBs (Fig 8). For the subcellular location, we tagged Snf7 with eYFP in a strain that also bore Nup49-mCherry as a NE reporter and Hta2-eCFP as a reporter of chromatin. Snf7-eYFP appeared concentrated in cytosolic spots (Fig 8a), probably reflecting endomembranes structures such as transport vesicles and/or endosomes as it has been reported before ^35^. Interestingly, single cell quantification showed that Snf7-eYFP abundance was rather variable in asynchronous cultures, and its levels increased after phleomycin treatment (Fig 8a,b). This increase in Snf7 levels was further confirmed by Western blot against a Snf7 tagged with the HA epitope (Fig 8c,d).

**Figure 8.**
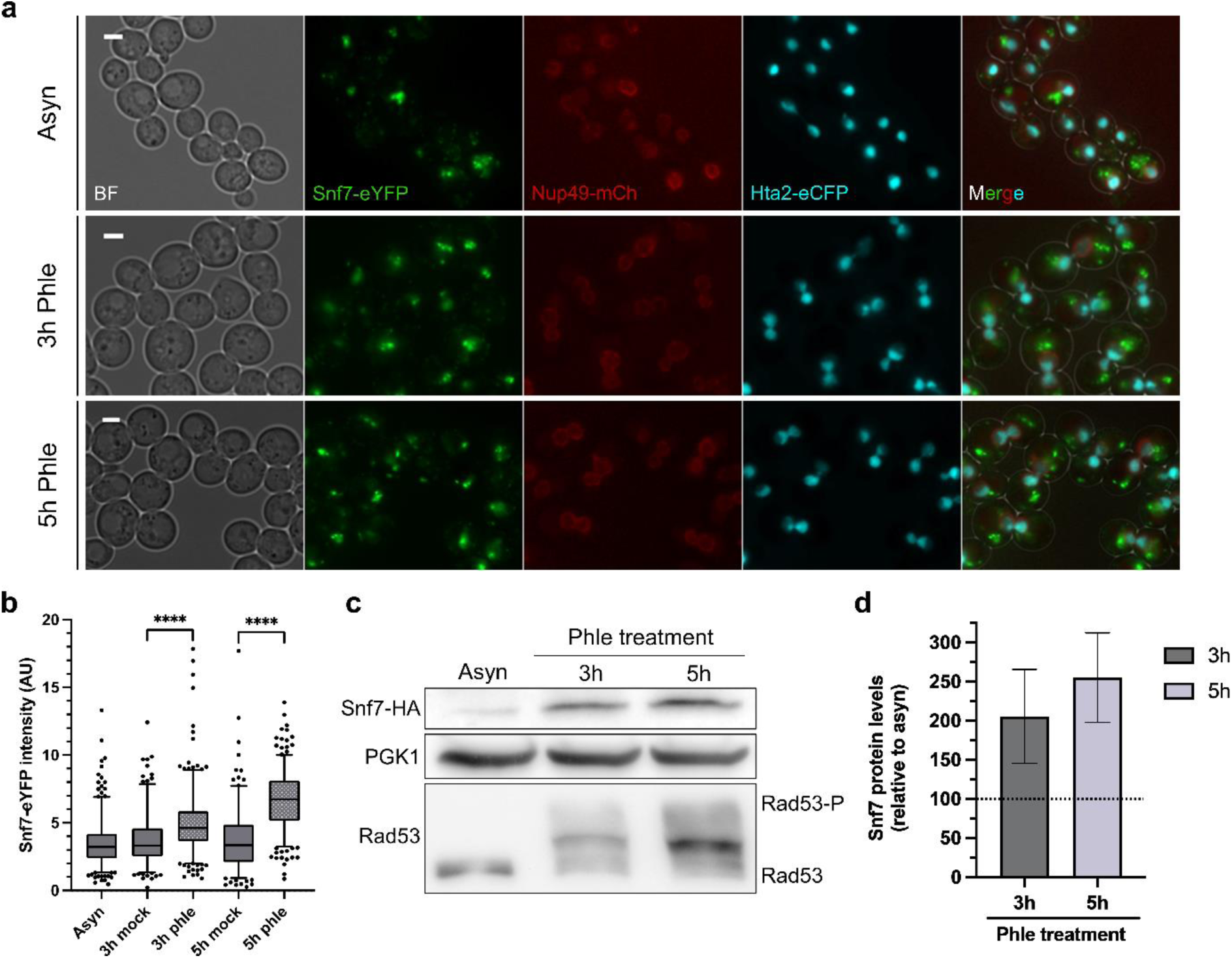
Snf7 levels increase after DSBs. **(a)** Representative micrographs of a strain carrying Snf7-eYFP, Nup49-mCherry and Hta2-eCFP. The first row corresponds to an asynchronous culture, while the subsequent two rows show the culture after generating DSBs with phleomycin for 3 hours and 5 hours, respectively. Note that Snf7-eYFP does not label the NE in any condition. **(b)** Single cell analysis of Snf7-eYFP abundance before and after DSBs. The “asyn” sample corresponds to the initial state of the overnight culture before it was split for the mock and phle treatments. **(c)** Western blot of Snf7-HA abundance before and after DSBs. Pgk1 is used as a housekeeping and Rad53 hyperphosphorylation is included to confirm DNA damage after phleomycin. **(d)** Levels of Snf7-HA after DSBs relative to the pre-DSB state. Note that single cell and population experiments confirm the increase in Snf7 levels after DSBs.

In conclusion, both Msc1 and Snf7 are not only important for a proper repair of DSBs in late-M but also in other phases of the cell cycle, in which DSBs elicit a G2/M arrest. In this arrest, NE and NPCs abnormalities are also seen in cells depleted from Msc1 and/or Snf7.

## Discussion

In *S. cerevisiae*, HR remains active despite sister chromatids being already segregated ^9,10^. This predicts that there is a cell cycle window in late mitosis (anaphase and telophase) in which HR-driven DSB repair could be highly mutagenic. However, it appears that yeast cells might still use the intact sister as a HR template by partly reversing segregation ^10^. The molecular players that reinforce HR with the sister in late-M are loosely defined. In a previous work, we identified the spindle machinery and the kinesin 5 Cin8 as important in ensuring partial regression. Since yeast performed a closed mitosis and sisters must transverse back the narrow bud neck during regression, we theorized that the NE ought to be important as well. Accordingly, we subsequently identified through proteomics the poorly-characterized NE protein Msc1 as a late-M player for HR and DSB repair ^21^. Msc1 levels increase after DSBs in late-M, and Δ*msc1* mutants delay DSB repair. Mechanistically, Msc1 appears to affect either the formation of the presynaptic filament or the homology search afterward. Regardless, Msc1 functions downstream of the DSB end resection and the establishment of the DNA damage checkpoint ^21^. Now, we herewith show that the Msc1 functional globular part faces the perinuclear space (lumen) (Fig 1), like its *S. pombe* orthologs ^22^. This fact implies that the action of Msc1 on HR must be indirect. In this sense, the morphological changes of the late-M nucleus in this mutant are remarkable (Fig 2, S2 and S3), and Msc1 appears to dynamically correct these aberrations in the WT nucleus (Fig 2e and S4). It is not lost on us that the most striking aberration, the partition/septation (inner over-compartimentalization) of the extended late-M nucleus (Fig 2b,c and S2), which also involves the compartmentalization of part of the segregated DNA (Fig 2d and S3), should retard or completely prevent the full mobilization of chromatids that ensues the DSB, resulting in the inability of the resected DSB ends to find the sister chromatid (Fig 3). This scenario would explain the late-M special role, which could stem from the need to recruit a sister template that is no longer aligned (Fig 9 for a model). In addition, the elongated and stressed late-M nucleus may have a greater need to protect the NE structure, with Msc1 being recruited to compromised areas to restore the hemisphericity of the two nuclear halves (Fig 2e and S4). This need may be less urgent in the more spherical G2/M nucleus, which is likely less prone to such severe septation, although we still observed this phenotype in a G2/M arrest elicited by DSBs over an asynchronous population (Fig 7). A seemingly obvious corollary of this model for late-M cells would be premature karyokinesis, which could indeed stem from the compartmentalization itself, but would also be expected based on the previous results obtained with SpLes1 and SpIsh1 ^13,14^. However, we found modest evidence of karyokinesis completion in Δ*msc1*, about 5-10% higher than the WT only (Fig 2a and S1). Even in extreme cases, nuclear compartments and bulges still seem connected to each other in the late-M nucleus (Fig 2c, S2 and S3; movie S1). Alternatively, the nuclear bridge could be affected in other ways that prevent sister chromatids from moving across it. For example, SpLes1 localizes to the bridge stalk before karyokinesis and corrals NPC at the midzone of the bridge ^13^. Thus, the Msc1 deficiency could also interfere with the movement of NPC-attached DSBs in their search for a template (Fig 8c). In support of this, we found signs of NPC mislocation in Δ*msc1* (Fig 6 and 7). In addition to defects in the mobilization of the whole chromatin, free circulation of the repair factors/factories could also be lessened. Indeed, physical interference with the location of repair factors has been suggested as the cause of DNA repair impairments in Hutchinson-Gilford progeria syndrome fibroblasts, which are also characterized by abnormal nuclei with lobes, invaginations and compartmentalization, creating unfavorable regions in the nucleus for DSB repair ^36^.

**Figure 9.**
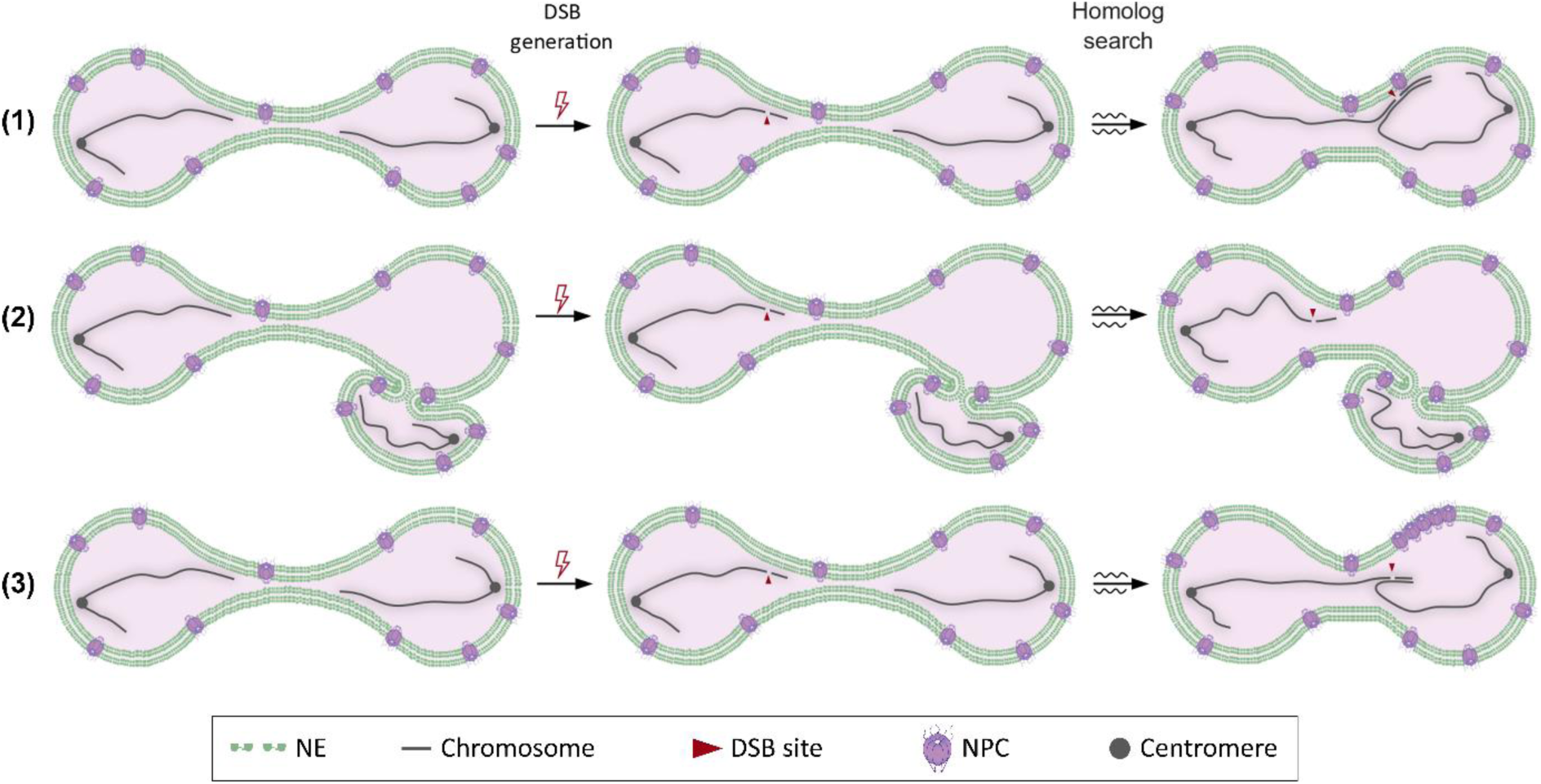
Models of how Msc1 and ESCRT-III may influence DSB repair in late-M. In late-M, the nucleus assumes an extended dumbbell shape with already segregated sister chromatids (a single pair of sisters is depicted for simplicity). (1) After a DSB occurs in one of the sisters, the extended nucleus becomes shorter and the bridge becomes thicker. The sisters also move closer and can coalesce for HR repair of the DSB. The broken ends can also get attached to the NE to facilitate retrograde events, and NPCs may participate in this attachment. (2) If segregated sisters are physically partitioned, as it occurs in *Δmsc1*, there are late-M cells in which sister chromatid approximation is challenged. A similar scenario may happen in NE invaginations in ESCRT-III mutants. (3) Alternatively, or in addition to partitions, NPC maldistribution and aggregation, which are observed in both mutants, could challenge template search.

ScMsc1 and SpLes1 are poorly-characterized NE proteins, in part due to the apparent lack of orthologs in higher eukaryotes. Nevertheless, SpLes1 is synthetically lethal with the highly conserved ESCRT-III complex ^16^. The fact that the absence of SpLes1 leads to transient NE ruptures, and the subsequent loss of the nucleo-cytoplasmic barrier in the extended late-M nucleus, suggests that the underlying nature of this negative genetic interaction resides in the need of sealing the ruptures by the ESCRT-III complex ^13,15^. It is fairly known that NE healing is one of the main cell functions of this versatile complex. For instance, HeLa cells that elongate their interphase nucleus to migrate through narrow spaces heavily rely on ESCRT-III to preserve NE integrity ^37,38^. Likewise, deregulation of ESCRT-III causes aberrant nuclear structures and micronuclei in HeLa and *D. melanogaster* cells ^38–40^, as well as invaginations of the INM in *C. elegans* ^41^. Because of the genetic interaction in *S. pombe*, and because the NE phenotypes resemble in part what we observed in the absence of Msc1, we decided to study the role of ESCRT-III in late-M DSB repair and its relationship with Msc1 (Fig 4-7 and S6-S14). Indeed, mutants for the core (Δ*snf7*) and regulatory (Δ*did4* and Δ*vps24*) components of ESCRT-III were hypersensitive to DSBs (Fig 4a, S6 and S7), and less capable of forming retrograde anaphase bridges after DSBs in late-M (Fig 4c). These mutants also rendered septated nuclei, but only at the G2/M arrest that ensues DSBs in an asynchronous population (Fig 7d). In late-M, septation was not seen and, instead, the major abnormal phenotype was a blurred NE (Fig 4d-g and S9, S10), which in turn was absent in G2/M. The most likely explanation for this difference is that septated nuclei do form in ESCRT-III mutants but, unlike in Δ*msc1*, they quickly collapse into the blurred nucleus in late-M. Indeed, high resolution airyscan microscopy suggests remnants of a septum in these blurred nuclei (Fig 4g). On the other hand, we looked into DSB repair and NE reshaping when both ESCRT-III and Msc1 were absent (Fig 5). We approached this by conditionally degrading ESCRT-III components in the Δ*msc1* background. We found no negative genetic interaction with this approach despite the levels of ESCRT-III dropping until becoming indetectable (Fig 5b, S11 and S12). Since the sensitivity to DSBs was higher in knockout mutants for ESCRT-III (compare Fig 4a and Fig 5c), we think that these conditional mutants have much less penetrance, which could also explain why they did not recreate in the Δ*msc1* background other NE phenotypes seen in the ESCRT-III knockouts (e.g., the blurred nuclei; Fig 5f). Regardless, there was synergism in DSB sensitivity and suppression of late-M retrograde events when Msc1 was absent and Snf7 diminished (Fig 5c-e and S12), which suggest a cooperation of both factors in DSB repair.

Another major phenotype of ESCRT-III mutants, also shared in the Δ*msc1* mutant, was the abnormal distribution of NPCs. This was seen in both the elongated NE of late-M cells and the more rounded NE of the G2/M arrest (Fig 6 and 7). It has been reported that ESCRT-III surveys for and/or eliminates non-functional NPCs, which otherwise aggregate into the so-called Storage of Improperly Assembled NPCs (SINC) compartment at the NE ^41^. Thus, the Msc1/ESCRT-III deficiency could also interfere with the axis that biochemically regulates the repair of DSBs via NPCs, which involves post-translational modifications of HR factors at the NPC and attachment of irreparable DSBs to the pores ^20,42–45^. In addition, the SINC can also cause NE breakdown. Whether through its direct NE healing activity or through SINC- or Msc1-mediated NE breakdown, ESCRT-III could also facilitate DSB repair by simply ensuring the nucleo-cytoplasmic barrier. In this alternative model, disruption of the NE as a barrier, rather than the NE partition or NPC malfunction, would leak in and out the cytoplasmic and nuclear proteins that undermine and ensure DNA repair, respectively. Future research ought to test these hypotheses. Remarkably, a strong association between major nuclear misshaping and SINC formation has been observed in other yeast mutants, including for processes as diverse as NPC assembly, chromatin remodeling, linkage of the nucleoskeleton to the cytoskeleton, and endoplasmic reticulum biology ^17,46–48^; and this association extends to both human and plant cells for at least some of these processes ^49,50^.

In conclusion, the NE protein Msc1 and the NE sealing complex ESCRT-III are critical for DSB repair in late-M and G2/M. Because the major phenotypes of their mutants affect the structure, integrity and shape of the NE, as well as the distribution of NPCs, we conclude that surveillance of the NE by these proteins is essential for DSB repair. To the best of our knowledge, this work presents for the first time the concept of active maintenance of the correct nuclear shape during DSB repair in yeast, and opens a new avenue of research in genome instability.

## Material and Methods

### Yeast strains and experimental conditions

Strains used in this work are listed in Table S1. Strain construction was undertaken through transformation, and mostly involved gene deletion and C-terminal tagging ^51^.

Cell cultures were grown overnight in air orbital incubators at 25 °C in YPD media (10 g·L^-1^ yeast extract, 20 g·L^-1^ peptone and 20 g·L^-1^ glucose). To arrest cells in late-M, log-phase asynchronous cultures were adjusted to OD_600_ ∼ 0.4 and the temperature was shifted to 34 °C for 3 h. In most experiments, the arrested culture was split into two subcultures: one subculture was treated with phleomycin (2 µg·mL^-1^; Sigma-Aldrich, P9564), and the one was left untreated (mock). In a few experiments, a third subculture was treated with β-estradiol (2 µM; Sigma-Aldrich, E8875) for the induction of HO endonuclease, which generates DSBs at the HO locus ^21,52^. Samples were collected at the arrest and 1 and 2 h after treatment. To synchronize cells in G2/M by the DNA damage checkpoint, phleomycin was added to the log-phase asynchronous population and the culture was held at 25°C for 3 h. In experiments with conditional degron variants for the auxin system (AID* tags), 5 mM 3-indol-acetic acid (IAA; Sigma-Aldrich, I2886) was added 1 h prior to adding phleomycin or β-estradiol.

### Western blots

Western blotting was carried out as reported before with minor modifications ^10,21^. Briefly, 5 ml samples were pelleted and fixed in 1 mL of 20% (w/v) trichloroacetic acid TCA. Cells were broken by vortexing for 3 min with ∼200 mg of glass beads. After centrifuging, pellets were resuspended in 150 µL of PAGE Laemmli Sample Buffer 1X (Bio-Rad, 1610747), Tris HCl 0.75M pH 8.0 and β-mercaptoethanol 2.5% (Sigma-Aldrich, M3148), and tubes were boiled at 95°C for 3 min and pelleted again. Total protein in the supernatant was quantified using a Qubit 4 Fluorometer (Thermo Fisher Scientific, Q33227). Proteins were resolved in 7.5% SDS-PAGE gels and transferred to PVFD membranes (Pall Corporation, PVM020C099). The membrane was stained with Ponceau S solution (PanReac AppliChem, A2935) as a loading reference.

The following antibodies were used for immunoblotting: The HA epitope was detected with a primary mouse monoclonal anti-HA (1:2,500; Sigma-Aldrich, H9658); the Myc epitope was detected with a primary mouse monoclonal anti-Myc (1:5,000; Sigma-Aldrich, M4439); the Pgk1 protein was recognized with a primary mouse monoclonal anti-Pgk1 (1:5,000; Thermo Fisher Scientific, 22C5D8), and Rad53 was recognized with a primary mouse monoclonal anti-Rad53 (1:2,500; Abcam, ab166859). A polyclonal goat anti-mouse conjugated to horseradish peroxidase (1:5,000 or 1:10,000; Promega, W4021) was used as the secondary antibody. Antibodies were diluted in 5% skimmed milk TBST (TBS pH 7.5 plus 0.1% Tween 20). Proteins were detected by using the ECL reagent (GE Healthcare, RPN2232) and visualized in a Vilber-Lourmat Fusion Solo S chamber. Protein bands were quantified using BioProfile Bio1D software (Vilber-Lourmat) and normalized to the housekeeping Pgk1.

### Microscopy

A Zeiss Axio Observer.Z1/7 was used for both wide field and confocal microscopy as reported before ^21^. The microscope was equipped for super-resolution confocal microscopy with live cell capability (LSM980 with Airyscan 2) together with the settings for wide field microscopy against CFP, YFP/GFP, and mCherry without crosstalk (Axiocam 702 sCMOS camera, the Colibri-7 LED excitation system, and narrow-band filter cubes). All images were obtained with a Plan-Apochromat 63x/NA 1.40 Oil M27 DIC objective. For each field, a stack of 10-20 z-focal planes (0.2-0.3 µm depth) was collected. In general, the images were taken from freshly harvested cells without further processing. For super-resolution images and 3D reconstructions, cells were first fixed with 3.7% w/v formaldehyde (Sigma-Aldrich, 47608) for 15 minutes. The Zen Blue (Zeiss) and Fiji-ImageJ (NIH) software were used for image processing and quantification.

### Growth curves, long-term viability spot assays, and short-term viability clonogenic assays

For real-time growth curves, log-phase overnight cultures were adjusted to an initial OD_600_ = 0.05 in YPD without or with phleomycin (2 µg·mL^-1^). Three replicates of each culture were aliquoted in a flat-bottomed 96-well plate and the real-time growth was measured in a Spark TECAN incubator by reading the OD_600_ every 15 minutes for 24 hours with shaking (96 rpm and 6mm of orbital amplitude). The mean of the three replicates was calculated to obtain the final growth curves. Two independent experiments were performed but only one is shown since the s.e.m was less than 0.1 OD_600_ for each time point.

For spot sensitivity assays cultures were grown exponentially and adjusted to an OD_600_ = 0.5 and then 5-fold serially diluted in YPD in 96-well plates. A 48-pin replica plater (Sigma-Aldrich, R2383) was used to spot ∼3 µL onto the corresponding plates, which were incubated at 25 °C for 3–4 days before taking photographs.

For clonogenic survival assays, log-phase asynchronous cultures were adjusted to OD_600_ = 0.4 before the corresponding treatment. After that, 100 µL of 4:10,000 dilutions were spread onto YPD plates. Viability was determined based on the number of colonies grown on the plates after 3 days at 25°C and normalized to the WT. The mock treatments yielded ∼500 colonies per plate.

### Data representation and statistics

Error bars in all graphs represent the standard error of the mean (s.e.m.) of independent biological replicates performed in different days. The number of replicates (n) is given in the figure legend. Graphpad Prism 10 was used for statistical tests. Differences between experimental data points were generally estimated using the Mann-Whitney U test, the unpaired t-test or ANOVA; the test used in each specific case is indicated in the figure caption.

In general, we used three types of graphs to represent the data: bar charts, XY line graphs and box plots. In box plots, the center line represents the medians, box limits represent the 25th and 75th percentile, the whiskers extend to the 5th and 95th percentiles, and the dots represent outliers.

## Data Availability

All data is contained within the manuscript and/or supplementary files.

## Acknowledgements

We would like to kindly thank Sue Jaspersen, Sarah Zanders, Rachel Helston and Lorraine Symington for strains and plasmids.

This research was funded by the Spanish Ministry of Science and Innovation (MCIN/AEI/10.13039/501100011033) through the grant PID2021-123716OB-I100 to F.M, which is co-funded by the EU-ERDF “A way of making Europe”. The Agencia Canaria de Investigación, Innovación y Sociedad de la Información (ACIISI) supported S. M-S. through a predoctoral fellowship (TESIS2020010028), co-funded by the EU-ESF+.

## Author contributions

S. M-S. performed all experiments shown in the main and supplementary figures and tables; constructed strains and plasmids; prepared the corresponding figures and tables; and gave critical insights as to the direction and development of the study.

F. M. supervised S.M-S, gave critical insights as to the direction and development of the study; was responsible for funding acquisition and project administration; and wrote the manuscript.

## References

1. Aparicio, T., Baer, R. & Gautier, J. DNA double-strand break repair pathway choice and cancer. DNA Repair (Amst). 19, 169–75 (2014).

2. Jeggo, P. A. & Löbrich, M. DNA double-strand breaks: their cellular and clinical impact? Oncogene 26, 7717–7719 (2007).

3. Trovesi, C., Manfrini, N., Falcettoni, M. & Longhese, M. P. Regulation of the DNA damage response by cyclin-dependent kinases. J. Mol. Biol. 425, 4756–66 (2013).

4. Kciuk, M., Gielecińska, A., Mujwar, S., Mojzych, M. & Kontek, R. Cyclin-dependent kinases in DNA damage response. Biochim. Biophys. acta. Rev. cancer 1877, 188716 (2022).

5. Scully, R., Panday, A., Elango, R. & Willis, N. A. DNA double-strand break repair-pathway choice in somatic mammalian cells. Nat. Rev. Mol. Cell Biol. 20, 698–714 (2019).

6. Thompson, R., Gatenby, R. & Sidi, S. How Cells Handle DNA Breaks during Mitosis: Detection, Signaling, Repair, and Fate Choice. Cells 8, 1049 (2019).

7. Audrey, A., de Haan, L., van Vugt, M. A. T. M. & de Boer, H. R. Processing DNA lesions during mitosis to prevent genomic instability. Biochem. Soc. Trans. 50, 1105–1118 (2022).

8. Machín, F. & Ayra-Plasencia, J. Are Anaphase Events Really Irreversible? The Endmost Stages of Cell Division and the Paradox of the DNA Double-Strand Break Repair. BioEssays 42, 2000021 (2020).

9. Ayra-Plasencia, J., Symington, L. & Machín, F. Cohesin still drives homologous recombination repair of DNA double-strand breaks in late mitosis. Elife 13, RP92706 (2024).

10. Ayra-Plasencia, J. & Machín, F. DNA double-strand breaks in telophase lead to coalescence between segregated sister chromatid loci. Nat. Commun. 10, 2862 (2019).

11. Boettcher, B. & Barral, Y. The cell biology of open and closed mitosis. Nucleus 4, 160–5 (2013).

12. Quevedo, O., García-Luis, J., Matos-Perdomo, E., Aragón, L. & Machín, F. Nondisjunction of a single chromosome leads to breakage and activation of DNA damage checkpoint in g2. PLoS Genet. 8, e1002509 (2012).

13. Dey, G. et al. Closed mitosis requires local disassembly of the nuclear envelope. Nature 585, 119–123 (2020).

14. Expósito-Serrano, M., Sánchez-Molina, A., Gallardo, P., Salas-Pino, S. & Daga, R. R. Selective Nuclear Pore Complex Removal Drives Nuclear Envelope Division in Fission Yeast. Curr. Biol. 30, 3212–3222.e2 (2020).

15. Stoten, C. L. & Carlton, J. G. ESCRT-dependent control of membrane remodelling during cell division. Semin. Cell Dev. Biol. 74, 50–65 (2018).

16. Frost, A. et al. Functional repurposing revealed by comparing S. pombe and S. cerevisiae genetic interactions. Cell 149, 1339–1352 (2012).

17. Webster, B. M., Colombi, P., Jäger, J. & Patrick Lusk, C. Surveillance of nuclear pore complex assembly by ESCRT-III/Vps4. Cell 159, 388–401 (2014).

18. Nagai, S. et al. Functional Targeting of DNA Damage to a Nuclear Pore–Associated SUMO-Dependent Ubiquitin Ligase. Science 322, 597 (2008).

19. Rodriguez-Berriguete, G. et al. Nucleoporin 54 contributes to homologous recombination repair and post-replicative DNA integrity. Nucleic Acids Res. 46, 7731–7746 (2018).

20. Gasser, S. M. & Stutz, F. SUMO in the regulation of DNA repair and transcription at nuclear pores. FEBS Lett. 597, 2833–2850 (2023).

21. Medina-Suárez, S., Ayra-Plasencia, J., Pérez-Martínez, L., Butter, F. & Machín, F. Msc1 is a nuclear envelope protein that reinforces DNA repair in late mitosis. iScience 27, 110250 (2024).

22. Asakawa, H., Hirano, Y., Shindo, T., Haraguchi, T. & Hiraoka, Y. Fission yeast Ish1 and Les1 interact with each other in the lumen of the nuclear envelope. Genes to Cells 27, 643–656 (2022).

23. Fedotova, A. A., Bonchuk, A. N., Mogila, V. A. & Georgiev, P. G. C2H2 Zinc Finger Proteins: The Largest but Poorly Explored Family of Higher Eukaryotic Transcription Factors. Acta Naturae 9, 47–58 (2017).

24. Smoyer, C. J. et al. Analysis of membrane proteins localizing to the inner nuclear envelope in living cells. J. Cell Biol. 215, 575–590 (2016).

25. D’Amours, D., Stegmeier, F. & Amon, A. Cdc14 and condensin control the dissolution of cohesin-independent chromosome linkages at repeated DNA. Cell 117, 455–69 (2004).

26. Machín, F., Torres-Rosell, J., Jarmuz, A. & Aragón, L. Spindle-independent condensation-mediated segregation of yeast ribosomal DNA in late anaphase. J. Cell Biol. 168, 209–19 (2005).

27. Matos-Perdomo, E., Santana-Sosa, S., Ayra-Plasencia, J., Medina-Suárez, S. & Machín, F. The vacuole shapes the nucleus and the ribosomal DNA loop during mitotic delays. Life Sci. alliance 5, e202101161 (2022).

28. Haber, J. E. Mating-Type Genes and MAT Switching in Saccharomyces cerevisiae. Genetics 191, 33–64 (2012).

29. Gnügge, R. & Symington, L. S. Efficient DNA double-strand break formation at single or multiple defined sites in the Saccharomyces cerevisiae genome. Nucleic Acids Res. 48, e115–e115 (2020).

30. Thaller, D. J. et al. An escrt-lem protein surveillance system is poised to directly monitor the nuclear envelope and nuclear transport system. Elife 8, 1–36 (2019).

31. Babst, M., Katzmann, D. J., Estepa-Sabal, E. J., Meerloo, T. & Emr, S. D. Escrt-III. Dev. Cell 3, 271–282 (2002).

32. Arii, J. et al. ESCRT-III mediates budding across the inner nuclear membrane and regulates its integrity. Nat. Commun. 9, 3379 (2018).

33. Nishimura, K., Fukagawa, T., Takisawa, H., Kakimoto, T. & Kanemaki, M. An auxin-based degron system for the rapid depletion of proteins in nonplant cells. Nat. Methods 6, 917–22 (2009).

34. Weinert, T. A. & Hartwell, L. H. The RAD9 gene controls the cell cycle response to DNA damage in Saccharomyces cerevisiae. Science 241, 317–22 (1988).

35. Babst, M. The Vps4p AAA ATPase regulates membrane association of a Vps protein complex required for normal endosome function. EMBO J. 17, 2982–2993 (1998).

36. Constantinescu, D., Csoka, A. B., Navara, C. S. & Schatten, G. P. Defective DSB repair correlates with abnormal nuclear morphology and is improved with FTI treatment in Hutchinson-Gilford progeria syndrome fibroblasts. Exp. Cell Res. 316, 2747–2759 (2010).

37. Denais, C. M. et al. Nuclear envelope rupture and repair during cancer cell migration. Science (80-.). 352, 353–358 (2016).

38. Vietri, M. et al. Unrestrained ESCRT-III drives micronuclear catastrophe and chromosome fragmentation. Nat. Cell Biol. 22, 856–867 (2020).

39. Willan, J. et al. ESCRT-III is necessary for the integrity of the nuclear envelope in micronuclei but is aberrant at ruptured micronuclear envelopes generating damage. Oncogenesis 8, 29 (2019).

40. Warecki, B., Ling, X., Bast, I. & Sullivan, W. ESCRT-III–mediated membrane fusion drives chromosome fragments through nuclear envelope channels. J. Cell Biol. 219, (2020).

41. Shankar, R., Lettman, M. M., Whisler, W., Frankel, E. B. & Audhya, A. The ESCRT machinery directs quality control over inner nuclear membrane architecture. Cell Rep. 38, 110263 (2022).

42. Mekhail, K., Seebacher, J., Gygi, S. P. & Moazed, D. Role for perinuclear chromosome tethering in maintenance of genome stability. Nature 456, 667–70 (2008).

43. Lemaître, C. et al. The nucleoporin 153, a novel factor in double-strand break repair and DNA damage response. Oncogene 31, 4803–9 (2012).

44. Géli, V. & Lisby, M. Recombinational DNA repair is regulated by compartmentalization of DNA lesions at the nuclear pore complex. Bioessays 37, 1287–92 (2015).

45. Loeillet, S. et al. Genetic network interactions among replication, repair and nuclear pore deficiencies in yeast. DNA Repair (Amst). 4, 459–68 (2005).

46. Titus, L. C., Dawson, T. R., Rexer, D. J., Ryan, K. J. & Wente, S. R. Members of the RSC Chromatin-Remodeling Complex Are Required for Maintaining Proper Nuclear Envelope Structure and Pore Complex Localization. Mol. Biol. Cell 21, 1072–1087 (2010).

47. Wente, S. R. & Blobel, G. A temperature-sensitive NUP116 null mutant forms a nuclear envelope seal over the yeast nuclear pore complex thereby blocking nucleocytoplasmic traffic. J. Cell Biol. 123, 275–84 (1993).

48. Liu, Q. et al. Functional association of Sun1 with nuclear pore complexes. J. Cell Biol. 178, 785–98 (2007).

49. Clever, M., Funakoshi, T., Mimura, Y., Takagi, M. & Imamoto, N. The nucleoporin ELYS/Mel28 regulates nuclear envelope subdomain formation in HeLa cells. Nucleus 3, 187–99 (2012).

50. Tamura, K. & Hara-Nishimura, I. Involvement of the nuclear pore complex in morphology of the plant nucleus. Nucleus 2, 168–72 (2011).

51. 51. Dunham, M., Gartenberg, M. & Brown, G. W. Methods in Yeast Genetics and Genomics, 2015 Edition: A CSHL Course Manual. (Cold Spring Harbor Laboratory Press, 2015).

52. Ottoz, D. S. M., Rudolf, F. & Stelling, J. Inducible, tightly regulated and growth condition-independent transcription factor in Saccharomyces cerevisiae. Nucleic Acids Res. 42, (2014).

