## Supplemental Figures and Tables for "Continuous nuclear envelope surveillance is required for DNA double-strand break repair"

### Supplemental Information

### SUPPLEMENTARY FIGURES

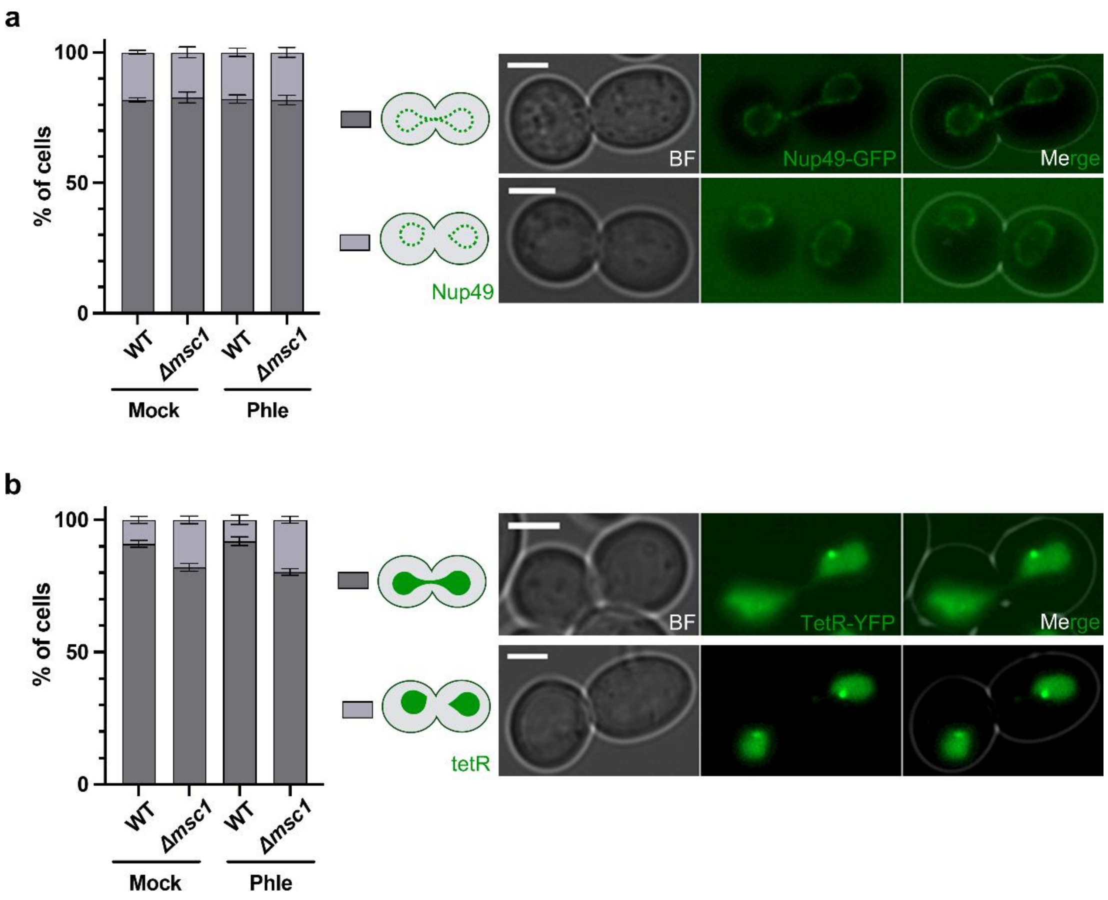


**Figure S1. Karyokinesis status in the WT and Δ*msc1* strains with and without DSB generation.** Related to Figure 2a. **(a)** Karyokinesis status as reported by the presence (gray bars) or absence (purple bars) of a Nup49-GFP NE bridge (mean ± s.e.m., n=3). Representative micrographs are shown on the right. **(b)** Karyokinesis status as reported by the presence (gray bars) or absence (purple bars) of a TetR-YFP nucleoplasmic bridge (mean ± s.e.m., n=2). Representative micrographs are shown on the right. Scale bars correspond to 3 µm. BF, bright field.

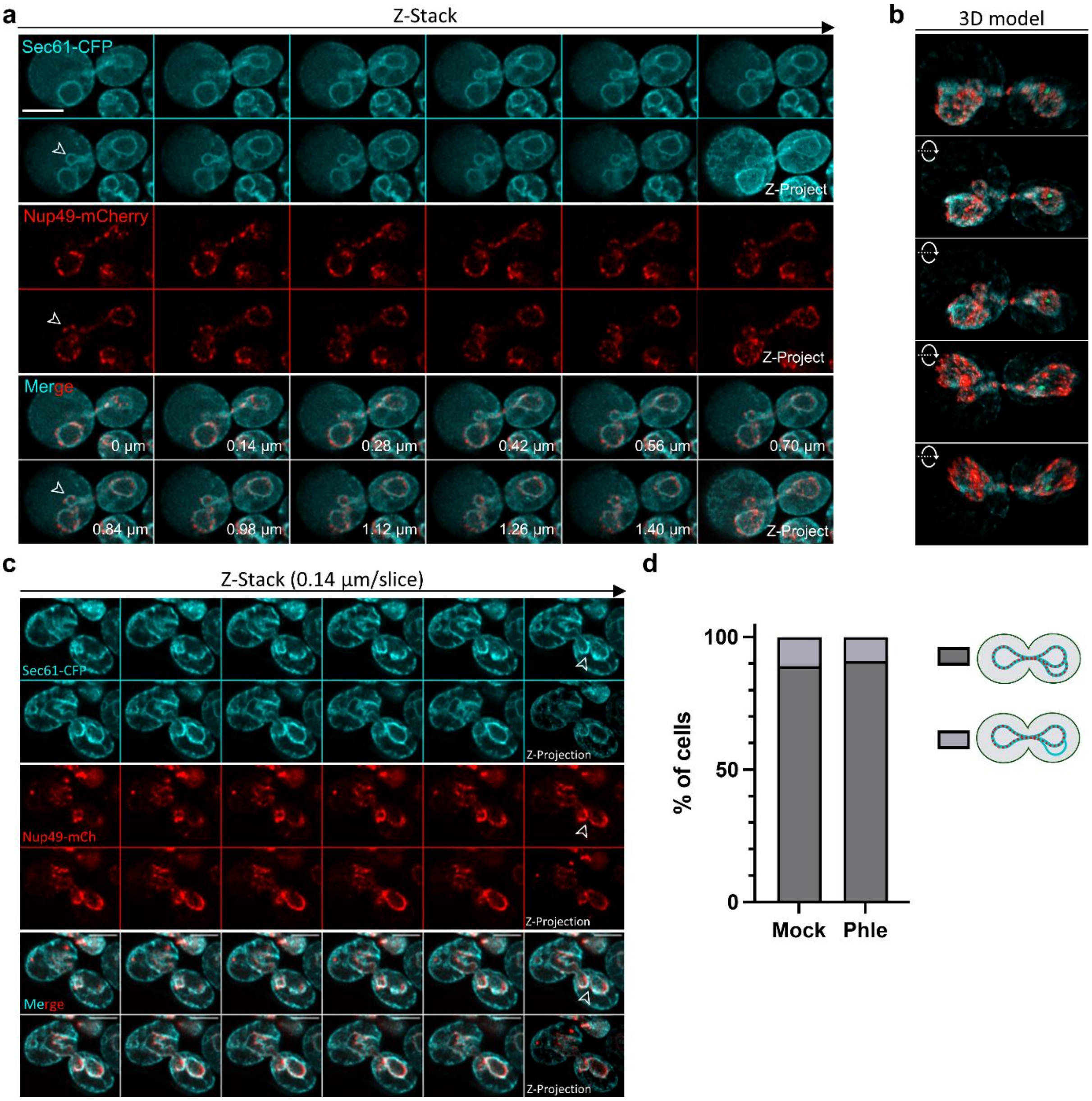


**Figure S2. High resolution of nuclear partitions in late-M *Δmsc1* cells.** Cells expressing both Sec61-eCFP and Nup49-mCherry were arrested in late-M and visualized by confocal airyscan 2 superresolution. **(a, b)** A full representation of the fixed cells shown in Figure 2c. **(a)** Z-series of that cell. Note that one of the segregating nuclear bodies presents a partition (indicated by the arrowhead). The NE septum contains NPCs. **(b)** 3D reconstruction of the same cell. Each micrograph represents a ~35º rotation on the X-axis. **(c)** Another example of a late-M *Δmsc1* cell. **(d)** Quantification of the presence of NPCs (Nup49) in the partitions observed with Sec61. NPCs are present in 90% of partitions.

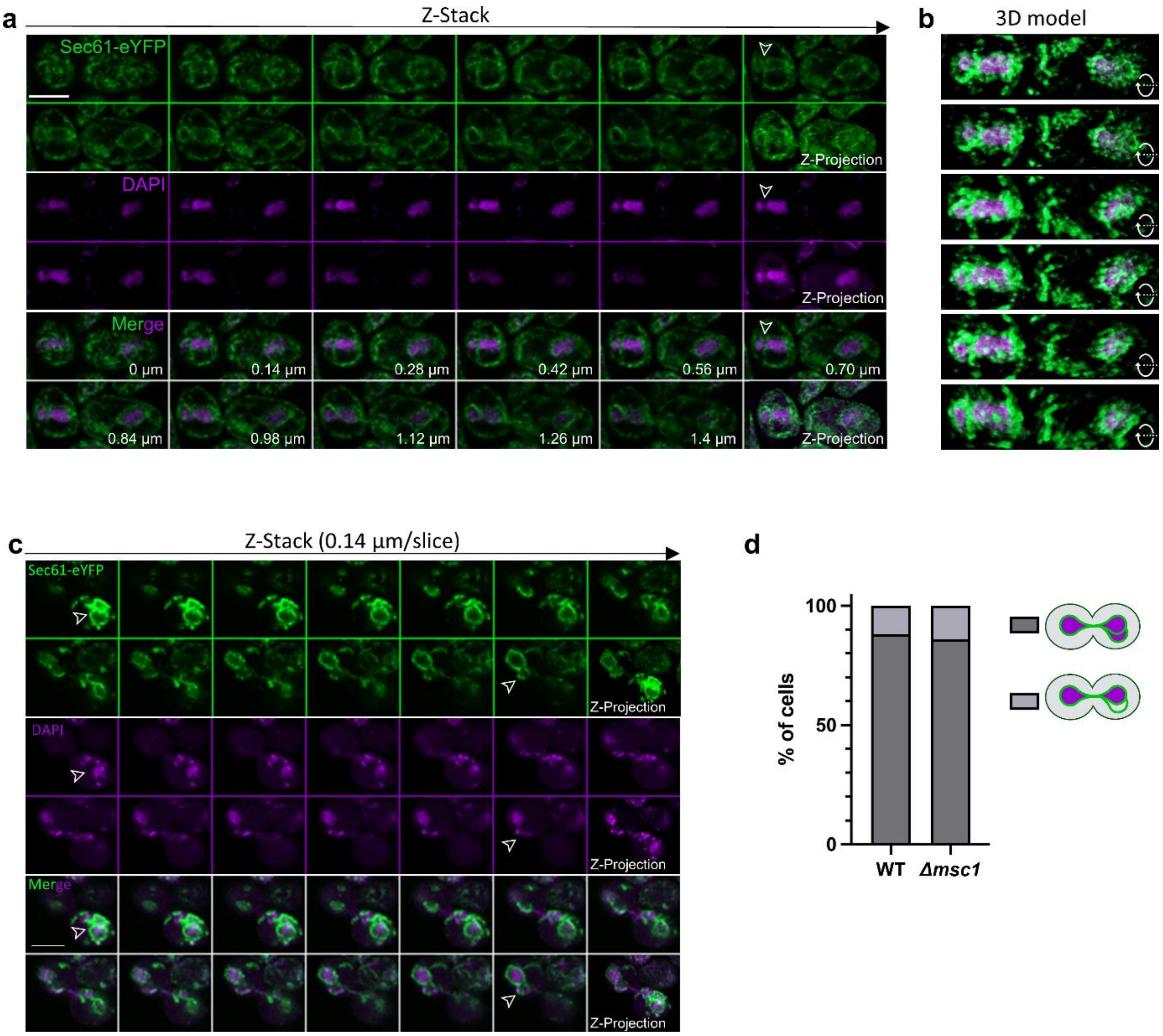


**Figure S3. Nuclear partitions contain DNA.** Cells expressing Sec61-eYFP were arrested in late-M, stained with DAPI, and visualized as in Figure S2. **(a, b)** A more complete representation of the fixed cells shown in Figure 2d. **(a)** Z-series of that cell. Note that the partition (indicated by the arrowhead) splits the nuclear DNA into two lobes. **(b)** 3D reconstruction of the same cell. Each micrograph represents a ~35° rotation on the X axis. **(c)** Another example of a late-M *Δmsc1* cell. **(d)** Quantification of the presence of split DNA masses in the partitions observed with Sec61. This occurs in 90% of partitions.

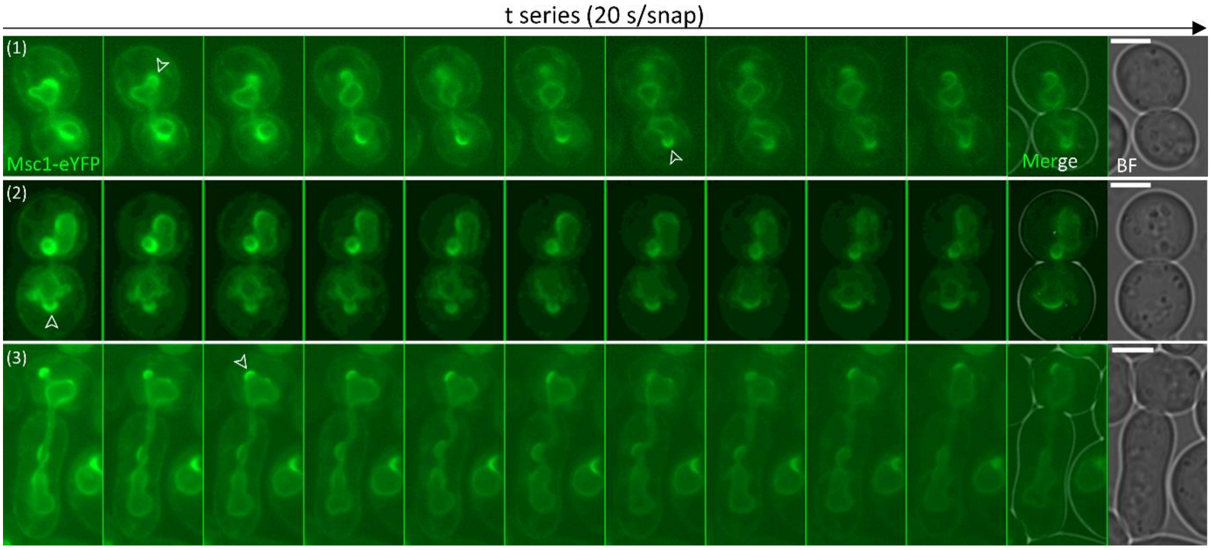


**Figure S4. Dynamics of Msc1 patches.** Related to Figure 2e. Short t-series of three examples of WT cells expressing Msc1-eYFP in late-M under phleomycin treatment. Areas of the NE with Msc1 patches exhibit greater plasticity and dynamics than the rest (indicated by arrowheads). Scale bars correspond to 3 µm. BF, bright field.

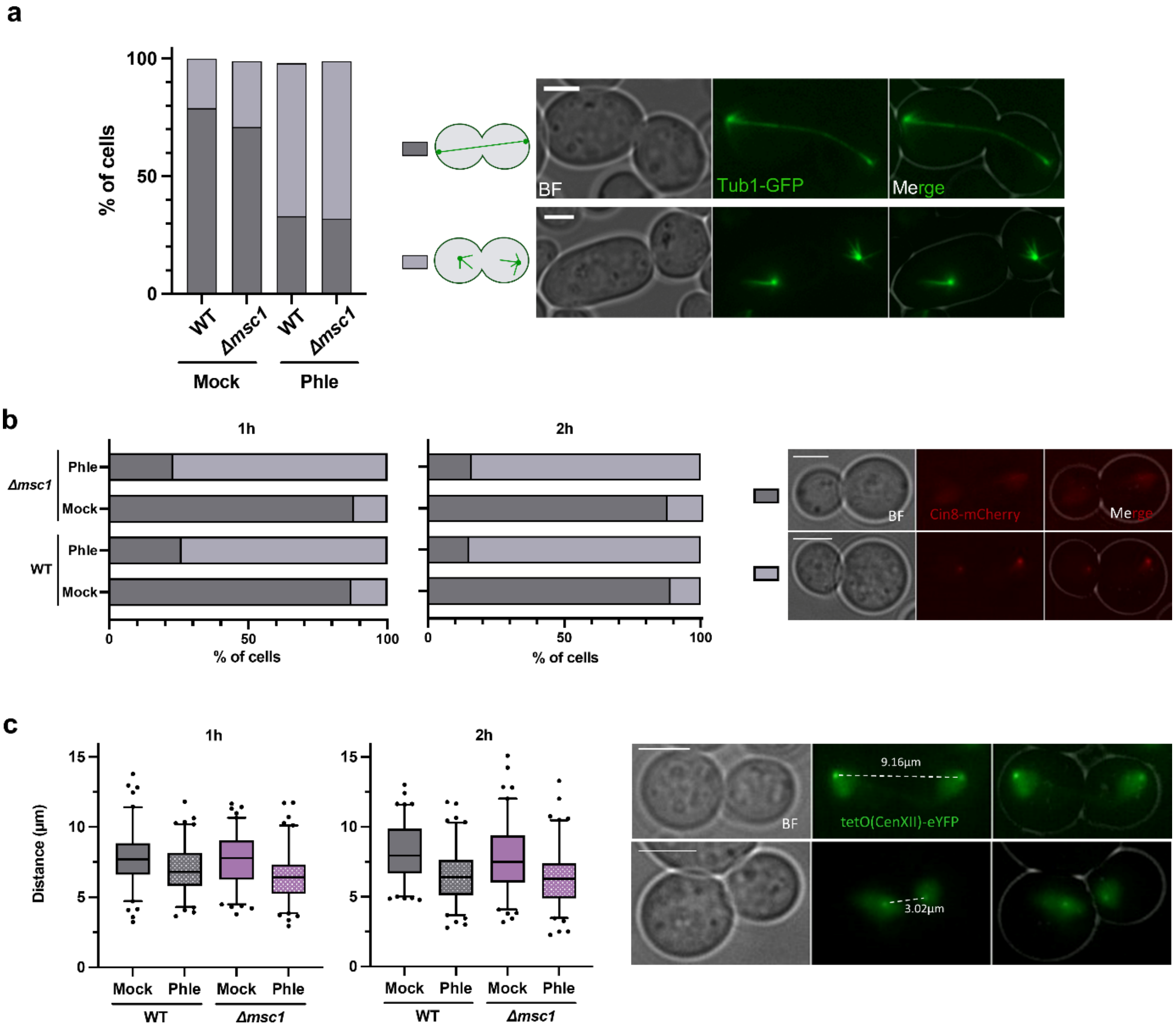


**Figure S5. Anaphase regression profiles of WT and Δ*msc1* after DSBs in late-M.** WT and Δ*msc1* strains carrying different reporters to assess anaphase progression were treated as in Figure S1. (**a**) Microtubule cytoskeleton after DSBs in late-M. Microtubules (as seen by GFP-Tub1) were categorized based on the presence or absence of the late anaphase spindle. Representative micrographs are shown on the right. (**b**) Cin8 relocalization after DSBs in late-M. Location of Cin8 was categorized into two groups: (i) all along the spindle and/or nucleoplasm (gray bars); and (ii) concentrated in two foci, presumably SPBs or kinetochores (purple bars). A representative from two independent experiments is shown. Representative micrographs are shown on the right. (**c**) Centromere approximation after DSBs in late-M. WT and Δ*msc1* strains carry the TetR-YFP and the *tetO* array integrated next to the cXII centromere (*tetO:194*). Distances between sister centromeres were box-plotted at the indicated time points. Note how the centromeres get closer in the same manner in the WT and the Δ*msc1* strains. On the right, representative late-M cells from the mock and phleomycin treatments (upper and lower panels, respectively). Scale bars correspond to 3 µm. BF, bright field.

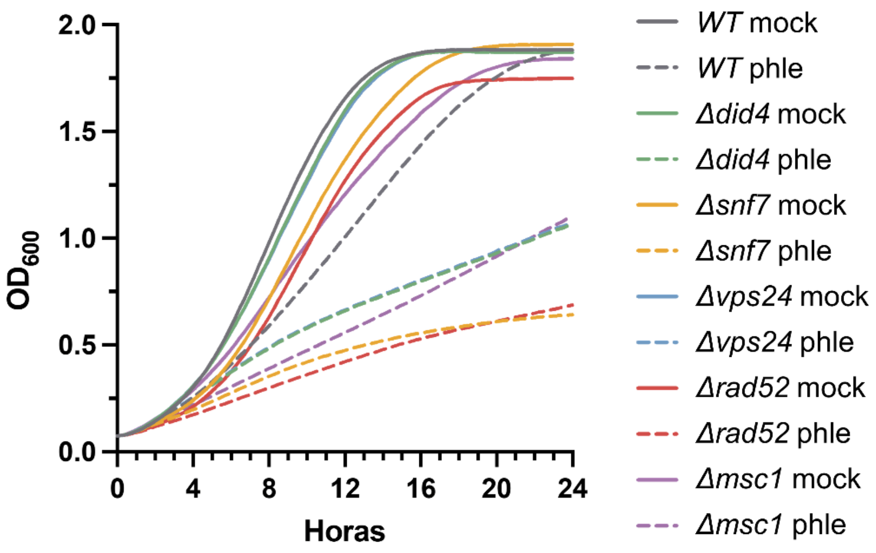


**Figure S6. ESCRT-III mutants are hypersensitive to DSBs in the YPH499 background.** Related to Figure 4a. Growth curves of the WT (YPH499 background) and isogenic knockout mutant strains for three ESCRT-III components (*did4*Δ, *vps24*Δ, and *snf7*Δ) without or with 2 µg/ml phleomycin. The isogenic *msc1*Δ and *rad52*Δ mutants were also included as controls. Note that ESCRT-III mutants are sensitive to phleomycin and that *snf7*Δ parallels *rad52*Δ.

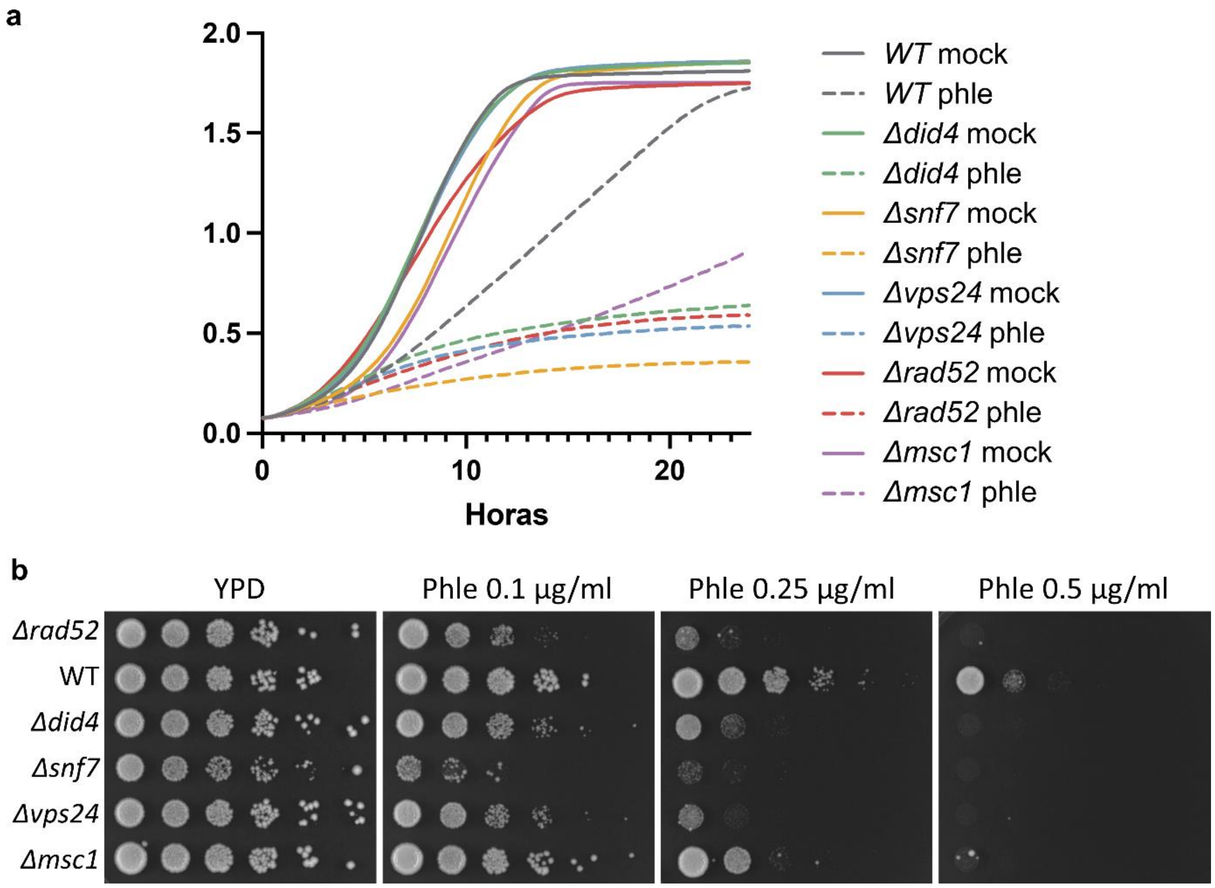


**Figure S7. ESCRT-III mutants are hypersensitive to DSBs in the BY4741 (S288C) background.** Related to Figure 4a. **(a)** Growth curves of the WT (BY4741) and isogenic knockout mutant strains for three ESCRT-III components (*did4*Δ, *vps24*Δ, and *snf7*Δ) without or with 2 µg/ml phleomycin. The isogenic *msc1*Δ and *rad52*Δ mutants were also included as controls. Note that ESCRT-III mutants are sensitive to phleomycin and that *snf7*Δ parallels *rad52*Δ. **(b)** Spot assays for the same strains under increasing concentrations of phleomycin. Note that ESCRT-III mutants are sensitive to phleomycin and that *snf7*Δ is even more sensitive than *rad52*Δ.

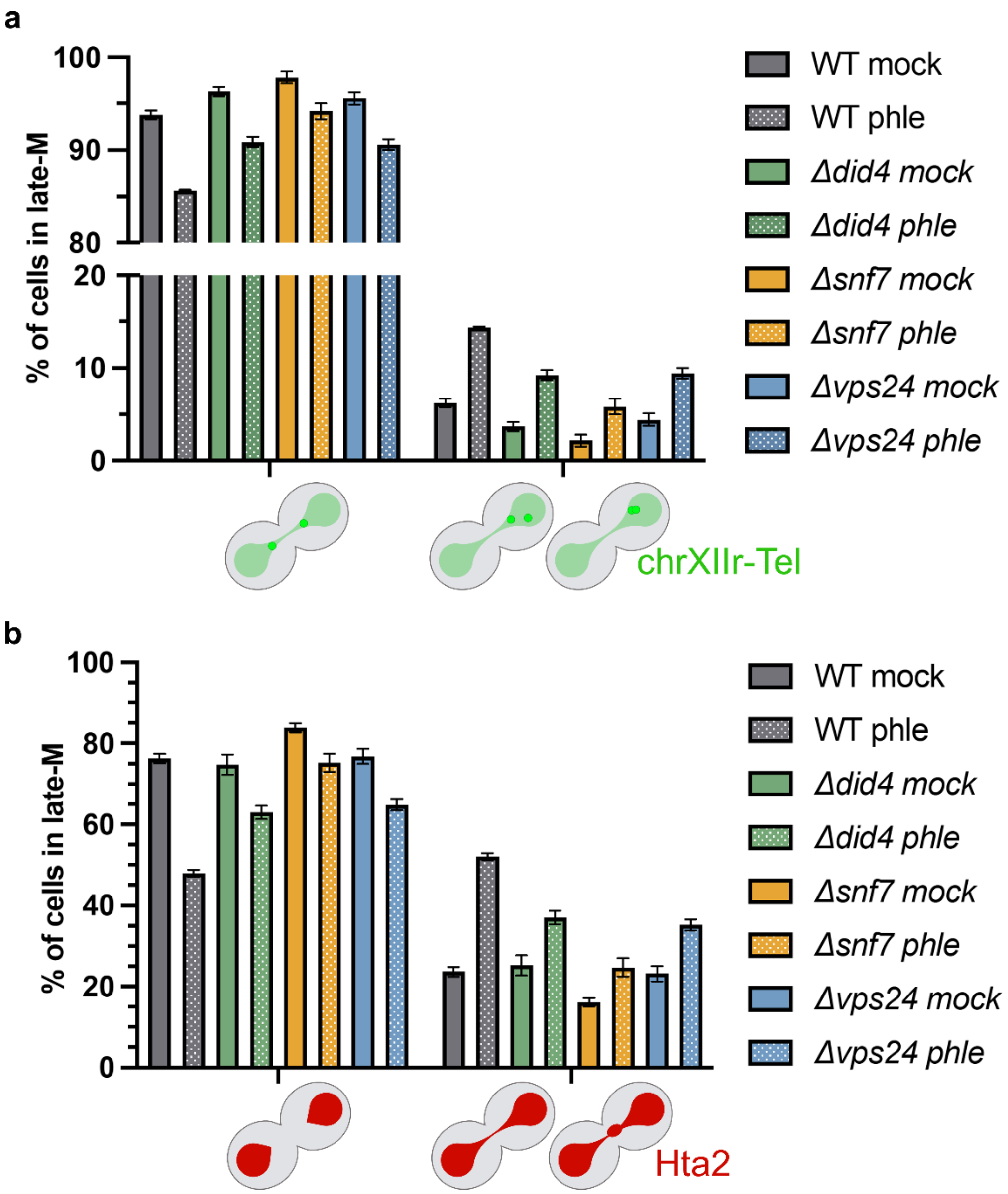


**Figure S8. Back migration and coalescence of sister loci in ESCRT-III mutants. (a)** WT and ESCRT-III mutants carrying a TetR-YFP/*tetOs* construct that labels the right telomere of chromosome XII (cXIIr-Tel) were treated as in Figure 3. Samples were taken 1 h after phle (or mock) addition, and late-M cells classified in two categories: segregated sister cXIIr-Tels and back migration of one sister cXIIr-Tel towards the other (mean ± s.e.m., n=3). **(b)** Like in panel (a) but with strains that label the bulk of chromatin with Hta2-mCherry (mean ± s.e.m., n=3). Late-M cells were classified in two categories again: without and with a chromatin bridge.

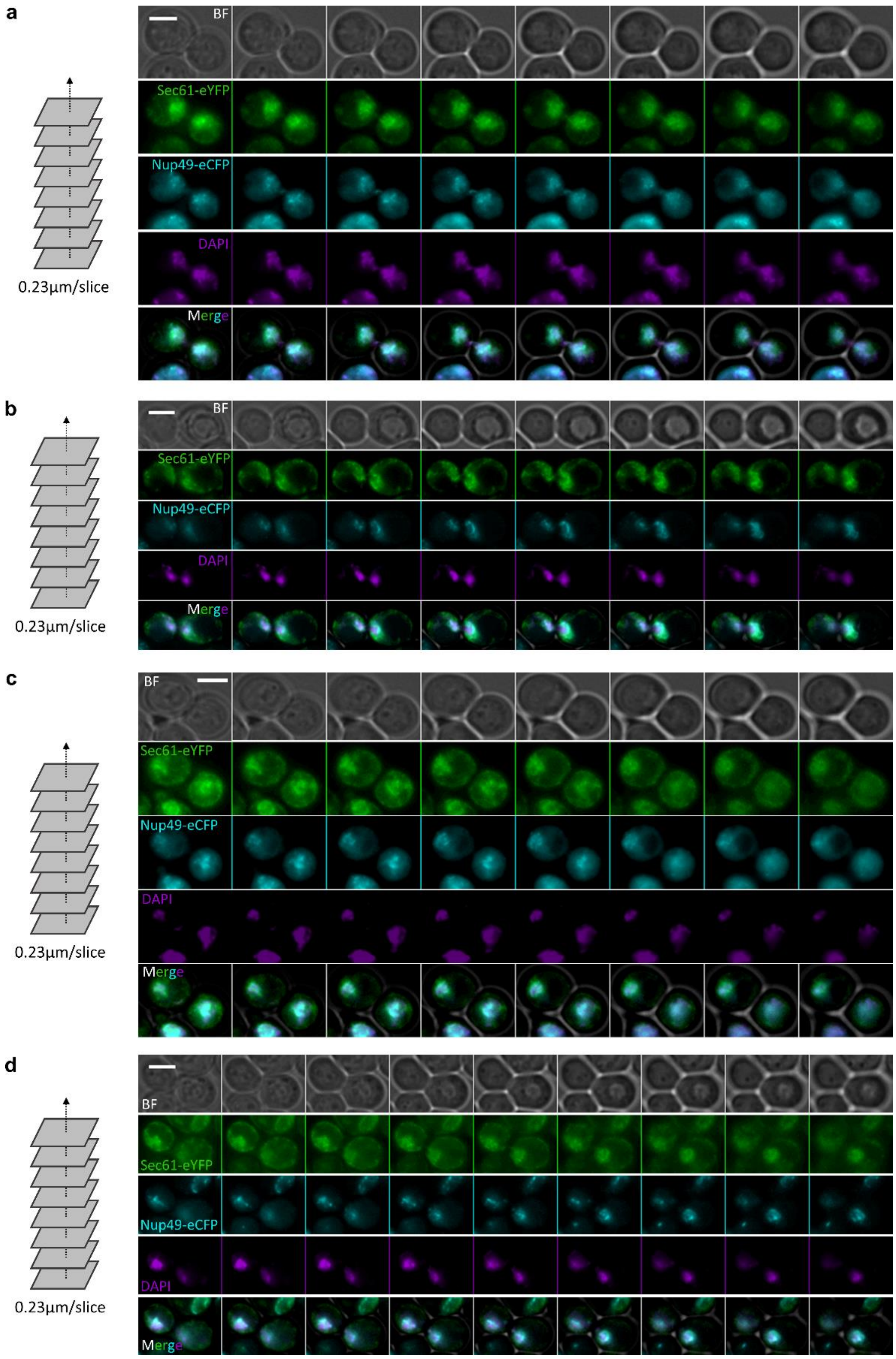


**Figure S9. Low resolution of blurred nuclear envelopes in ESCRT-III mutants.** Related to Figure 4d. Four examples of late-M *snf7*Δ Sec61-eCFP Nup49-mCherry cells with the blurred NE category (category "c"). Cells were visualized by epifluorescence microscopy (individual z slices shown) and the nuclear mass was stained by DAPI. All examples correspond to mock (no phleomycin) subcultures. Note that it is difficult, if not impossible, to outline the NE by the NE markers, even taking the DAPI as a reference. It is not only that NPCs (Nup49) appear misdistributed, but also Sec61. Scale bars correspond to 3 µm. BF, bright field.

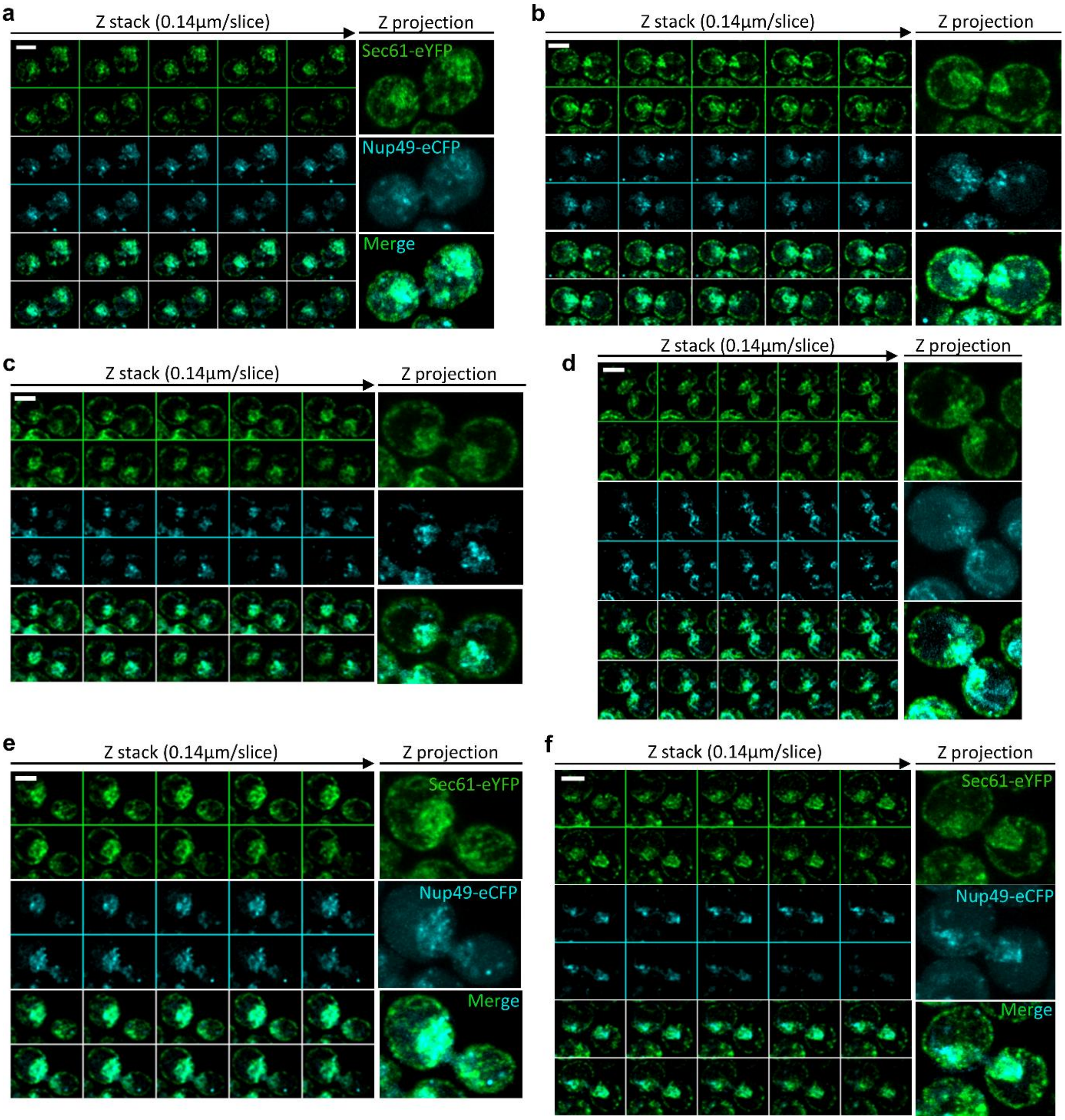


**Figure S10. High resolution of blurred nuclear envelopes in ESCRT-III mutants.** Related to Figure 4g. Six examples of late-M *snf7*Δ Sec61-eCFP Nup49-mCherry cells with the blurred NE category (category "c"). Cells were visualized by confocal airyscan 2 superresolution (individual z slices shown together with the 2D projection). All examples correspond to mock (no phleomycin) subcultures. Note that NE markers appear spotted all over the nuclear area, suggesting that the NE is not defined. Scale bars correspond to 3 µm.

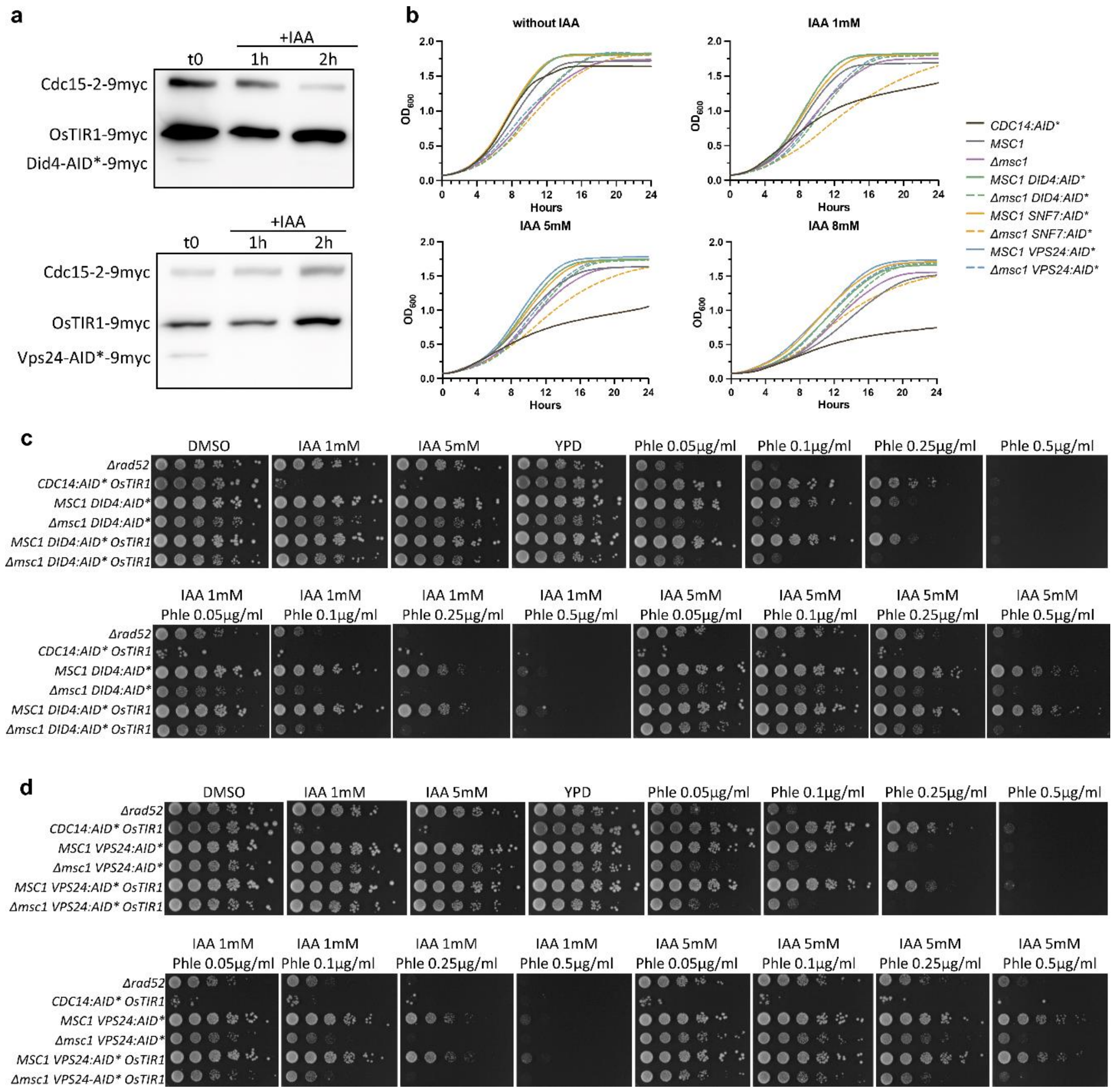


**Figure S11. Synergism profiles of Msc1 and ESCRT-III in DSB sensitivity.** Related to Figure 5. **(a)** Western blot to show that the core ESCRT-III components Did4 and Vps24 tagged with the AID* degron are degraded upon auxin addition (+IAA). The blot was probed with an anti-myc antibody, which recognizes three tagged proteins in these strains; Did4- or Vps24-AID*-9myc proper, Cdc15-2-9myc and OsTIR1-9myc. The latter two served as loading controls. (**b**) Growth curves of the WT (*MSC1*), *msc1*Δ and isogenic AID*-tagged derivative strains for three ESCRT-III components (Snf7, Did4 and Vps24) under increasing concentrations of IAA. The *CDC14:AID* OsTIR1* strain was included as a positive control for IAA. (**c**) Spot assay against combinations of IAA and Phle of *did4:aid* Δmsc1* double mutants. The *CDC14:AID* OsTIR1* strain was included as a control for IAA. The *Δrad52* strain was included as a control for phle. *MSC1* and just *did4:aid** (no OsTIR1) strains were also included as a reference for the putative genetic interaction and to assess the effect of Did4 C-terminal tagging, respectively. (**d**) Spot assay against combinations of IAA and Phle of *vps24:aid* Δmsc1* double mutants. The *CDC14:AID* OsTIR1* strain was included as a control for IAA. The *Δrad52* strain was included as a control for phle. *MSC1* and just *vps24:aid** (no OsTIR1) strains were also included as a reference for the putative genetic interaction and to assess the effect of Vps24 C-terminal tagging, respectively.

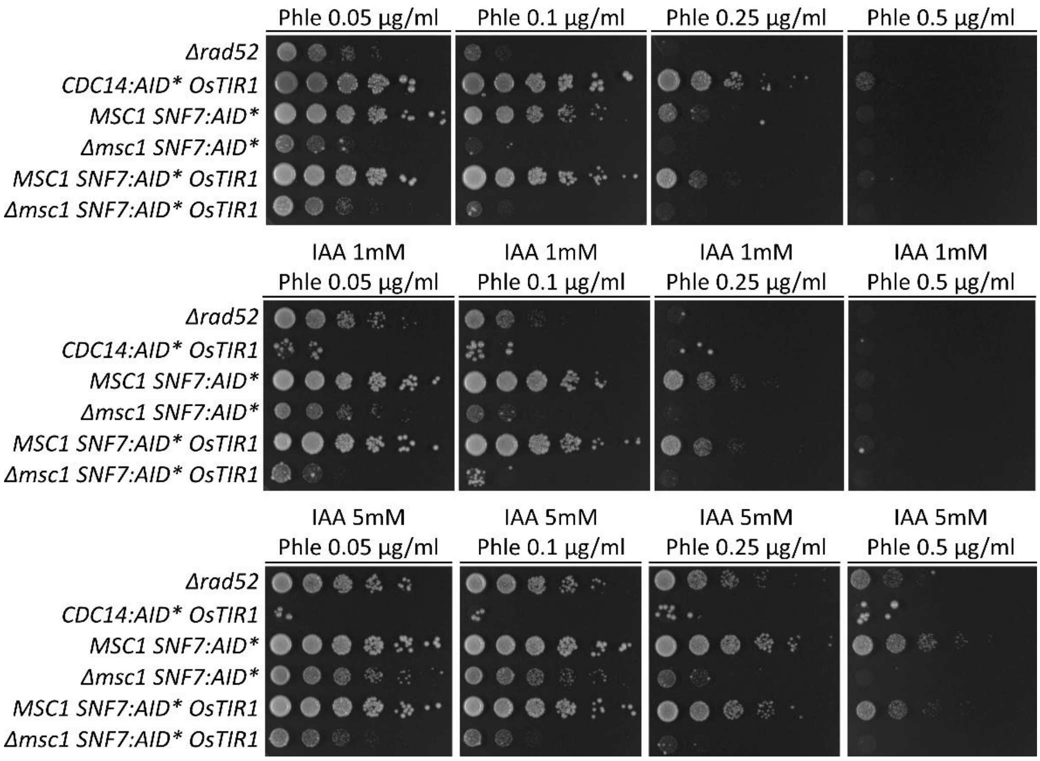


**Figure S12. Synergism profiles of Msc1 and Snf7 in DSB sensitivity.** Related to Figure 5c. Spot assay against combinations of IAA and Phle of *snf7:aid* Δmsc1* double mutants. The *CDC14:AID* OsTIR1* strain was included as a positive control for IAA. The *Δrad52* strain was included as a positive control for phle. *MSC1* and just *snf7:aid** (no OsTIR1) strains were also included as a reference for the putative genetic interaction and to assess the effect of Snf7 C-terminal tagging, respectively.

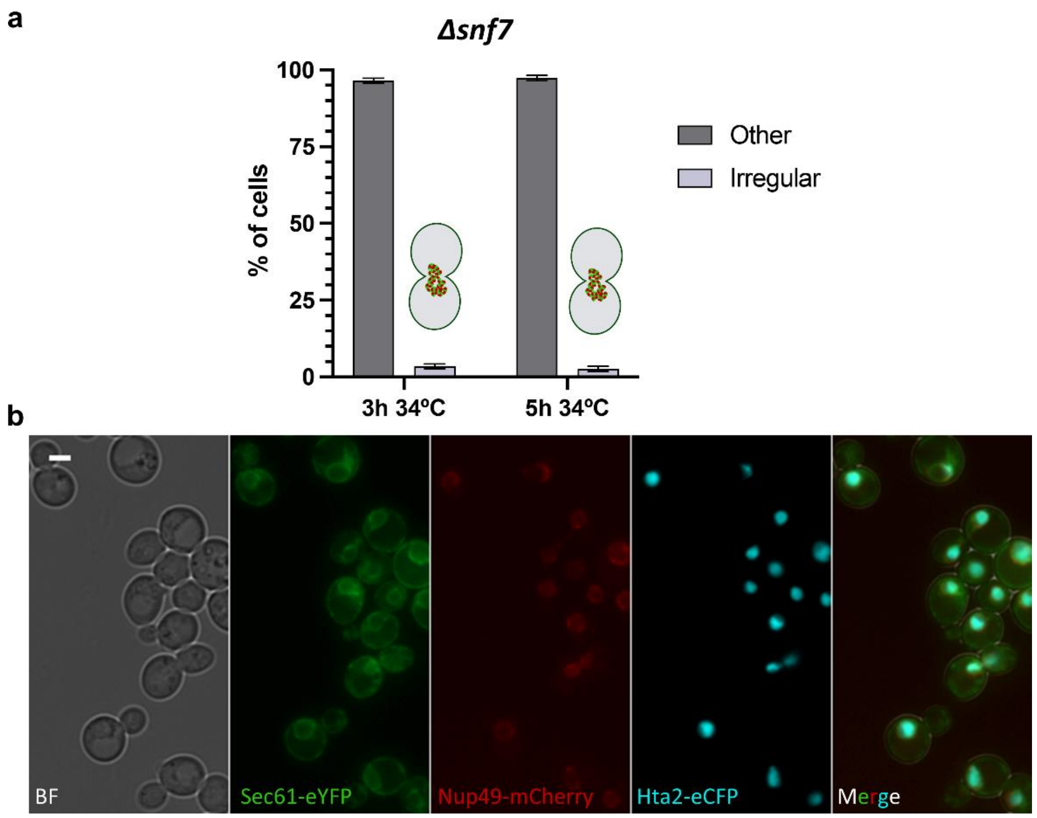


**Figure S13. Effect of the temperature in the nuclear envelope morphology in the *snf7*Δ mutant.** The *snf7*Δ strain expresses the NE markers Sec61-eYFP and Nup49-mCherry as well as chromatin marker Hta2-eCFP. The strain was grown in YPD at 25 ºC to log-phase and shifted to 34 ºC for the indicated time. (**a**) Quantification of the blurred NE after the shift. (**b**) Representative microscopy field of the asynchronous culture at 34 ºC. Scale bars correspond to 3 µm. BF, bright field.

**SUPPLEMENTARY TABLES**

**Table S1. Strains used in this work.**

| Name | Genotype[1] | Origin | Figure |
|---|---|---|---|
| FM2995 | *MATα can1Δ::STE2p:SpHIS5 lyp1Δ his3Δ1 leu2Δ0 ura3Δ0 met15Δ0 LYS2; [NOP1p:GFP11:mCherry:PUS1::LEU2 (pSJ1321)]; MSC1:yGFP1–10::CaURA3MX* | This study | 1c |
| FM2996 | *MATα can1Δ::STE2p:SpHIS5 lyp1Δ his3Δ1 leu2Δ0 ura3Δ0 met15Δ0 LYS2; MSC1:yGFP1–10::CaURA3MX* | This study | 1c |
| FM2997 | *MATα can1Δ::STE2p:SpHIS5 lyp1Δ his3Δ1 leu2Δ0 ura3Δ0 met15Δ0 LYS2; [NOP1p:mCherry:SCS2TM:GFP11::LEU2 (pSJ1602)]; MSC1:yGFP1–10::CaURA3MX* | This Study | 1c |
| FM2999 | *MATα can1Δ::STE2pr-SpHIS5 lyp1Δ his3Δ1 leu2Δ0 ura3Δ0 met15Δ0 LYS2; [NOP1p:GFP11:mCherry:SCS2TM::LEU2 (pSJ1568)]; MSC1:yGFP1–10::CaURA3MX* | This Study | 1c |
| FM2942 | *MATa bar1Δ leu2-3,112 ura3-52 his3-Δ200 trp1-Δ63 ade2-1 lys2-801; cdc15-2:9myc::Hph; HMLα Δhmr::HIS3MX; leu2-3::LexA-TF-PlexOp:HO::LEU2; SEC61:eYFP::KanMX* | This Study | 2a,b; 4a; S6 |
| FM2974 | *MATa bar1Δ leu2-3,112 ura3-52 his3-Δ200 trp1-Δ63 ade2-1 lys2-801; cdc15-2:9myc::Hph; HMLα Δhmr::HIS3MX; leu2-3::LexA-TF-PlexOp:HO::LEU2; Δmsc1::NatNT2; SEC61:eYFP::KanMX* | This Study | 2a,b,d; 4a; 7a-c; S3; S6 |
| FM2976 | *MATa bar1Δ leu2-3,112 ura3-52 his3-Δ200 trp1-Δ63 ade2-1 lys2-801; ade2-1:TetR:YFP::ADE2; tetO(5.6Kb)::1061Kb-ChrXII::HIS3; cdc15-2:9myc::Hph; NUP49:mCherry::KanMX6; Δmsc1::NatNT2; SEC61:CFP::TRP1* | This Study | 2c; 7d-e; S2 |
| FM2831 | *MATa bar1Δ leu2-3,112 ura3-52 his3-Δ200 trp1-Δ63 ade2-1 lys2-801; cdc15-2:9myc::Hph; HMLα Δhmr::HIS3MX; leu2-3::LexA-TF-PlexOp:HO::LEU2; MSC1:eYFP::KanMX* | This study | 2e; S4 |
| FM588 | *MATa bar1Δ leu2-3,112 ura3-52 his3-Δ200 trp1-Δ63 ade2-1 lys2-801; ade2-1::TetR:YFP::ADE2; tetO(5.6Kb)::1061Kb-ChrXII::HIS3; cdc15-2:9myc::HphNT1* | F. Machín lab | 3a-c; 4b; S1b; S8a; S10a; S11b |
| FM2807 | FM588; *Δmsc1::NatNT2* | This Study | 3a-c; S1b; S11b |
| FM2354 | *Mata bar1Δ leu2-3,112 ura3-52 his3-Δ200 trp1-Δ63 ade2-1 lys2-801; ade2-1::TetR:YFP::ADE2; tetO(5.6Kb)::194Kb-ChrXII::HIS3; cdc15-2:9myc::HphNT1; HTA2:mCHERRY::KanMX6* | F. Machín lab | 3d-f; 4c; S8b; S10b |

| | | | |
|---|---|---|---|
| FM3127 | FM2354; *Δmsc1::NatNT2* | This study | 3d-f |
| FM82 | *MATa bar1Δ leu2-3,112 ura3-52 his3-Δ200 trp1-Δ63 ade2-1 lys2-801; Δrad52::kanMX4* | Machín lab | 4a, 5b,c; S6 S11c,d; S12 |
| FM3129 | FM2942; *Δdid4::NatNT2* | This study | 4a; S6 |
| FM3130 | FM2942; *ΔSnf7::NatNT2* | This study | 4a; S6 |
| FM3131 | FM2942; *ΔVps24::NatNT2* | This study | 4a; S6 |
| FM3139 | FM588; *Δdid4::NatNT2* | This study | 4b; S8a; S1 |
| FM3142 | FM588; *Δsnf7::NatNT2* | This study | 4b; S8a; S10a |
| FM3145 | FM588; *Δvps24::NatNT2* | This study | 4b; S8a; S10a |
| FM3140 | FM2354; *Δdid4::NatNT2* | This study | 4c; S8b; S10b |
| FM3144 | FM2354; *Δsnf7::NatNT2* | This study | 4c; S8b; S10b |
| FM3147 | FM2354; *Δvps24::NatNT2* | This study | 4c; S8b; S10b |
| FM3149 | FM2942; *NUP49-eCFP::TRP1* | This study | 4d,e; 6a,b; 7 |
| FM3151 | FM3129; *NUP49-eCFP::TRP1* | This study | 4d,e; 6a,b; 7 |
| FM3152 | FM3130; *NUP49-eCFP::TRP1* | This study | 4d-g; 6a,b; 7; S9; S10 |
| FM3162 | FM3131; *NUP49-eCFP::TRP1* | This study | 4d,e; 6a,b; 7 |
| FM3170 | FM588; *SNF7:AID*:9myc::KanMX* | This study | 5b,c; S12 |
| FM3172 | FM2807; *SNF7:AID*:9myc::KanMX* | This study | 5b,c; S12 |
| FM3188 | FM3170; *ura3-52::ADH1p:OsTIR1:9myc::URA3* | This study | 5a-c; S11b; S12 |
| FM3189 | FM3172; *ura3-52::ADH1p:OsTIR1:9myc::URA3* | This study | 5b-d; S11b; |

| | | | |
|---|---|---|---|
| | | | S12 |
| FM1391 | *Mata ura3-52::ADH1p:OsTIR1:9myc::URA3 ade2-1 his3-11,15 leu2-3,112 trp1-1 can1-100; NET1:GFP::LEU2; cdc14:aid::KanMX* | F. Machín lab | 5b,c; S11b-d; S12 |
| FM3198 | FM3189; *HTA2:eCFP::TRP1* | This study | 5e |
| FM3202 | FM3189; *SEC61:eCFP::TRP1* | This study | 5f |
| FM3200 | FM3189; *NUP49:eCFP::TRP1* | This study | 6c |
| FM3194 | *MATa bar1Δ leu2-3,112 ura3-52 his3-Δ200 trp1-Δ63 ade2-1 lys2-801; NUP49:mCherry::NatNT2; HTA2:eCFP::TRP1; SNF7::eYFP::HphNT1* | This study | 8a,b |
| FM3246 | *MATa bar1Δ leu2-3,112 ura3-52 his3-Δ200 trp1-Δ63 ade2-1 lys2-801; NUP49:mCherry::NatNT2; HTA2:eCFP::TRP1;SNF7:6HA::HIS3* | This study | 8c,d |
| FM2947 | *MATa bar1Δ leu2-3,112 ura3-52 his3-Δ200 trp1-Δ63 ade2-1 lys2-801; cdc15-2:9myc::Hph; HMLα Δhmr::HIS3MX; leu2-3::LexA-TF-PlexOp:HO::LEU2*; *RAD52:mCherry::NatNT2; NUP49:eGFP::TRP1* | This study | S1a |
| FM2954 | *MATa bar1Δ leu2-3,112 ura3-52 his3-Δ200 trp1-Δ63 ade2-1 lys2-801; cdc15-2:9myc::Hph; HMLα Δhmr::HIS3MX; leu2-3::LexA-TF-PlexOp:HO::LEU2*; *Δmsc1::NatNT2*; *RAD52:mCherry::KanMX; NUP49:eGFP::TRP1* | This study | S1a |
| FM2381 | *MATa bar1Δ leu2-3,112 ura3-52 his3-Δ200 trp1-Δ63 ade2-1 lys2-801; GFP:TUB1::URA3; cdc15-2:9myc::HphNT1* | F. Machín lab | S5a |
| FM2789 | FM2381; *Δmsc1::NatNT2* | This Study | S5a |
| FM2317 | *MATa bar1Δ leu2-3,112 ura3-52 his3-Δ200 trp1-Δ63 ade2-1 lys2-801; ade2-1::TetR:YFP::ADE2; tetO(5.6Kb)::194Kb-ChrXII::HIS3; cdc15-2:9myc::HphNT1; CIN8:mCherry::KanMX6* | F. Machín lab | S5b,c |
| FM2804 | FM2317; *Δmsc1::NatNT2* | This study | S5b,c |
| FM632 | *MATa his3Δ1 leu2Δ0 met15Δ0 ura3Δ0 Δrad52::kanMX4* | Euroscarf | S7 |
| FM23 | *MATa his3Δ1 leu2Δ0 met15Δ0 ura3Δ0* | Euroscarf | S7 |
| FM3002 | FM23; *Δdid4::HphNT1* | This study | S7 |
| FM3004 | FM23; *Δsnf7::NatNT2* | This study | S7 |
| FM3003 | FM23; *Δvps24::NatNT2* | This study | S7 |
| FM2865 | *MATα his3Δ1 leu2Δ0 lys2Δ0 ura3Δ0; Δmsc1:NatNT2* | This study | S7 |
| FM3166 | FM588; *DID4:AID*:9myc::KanMX* | This study | S11c |

| | | | |
|---|---|---|---|
| FM3168 | FM2807; *DID4:AID*:9myc::KanMX* | This study | S11c |
| FM3186 | FM3166; *ADH1p:OsTIR1-9myc::URA3* | This study | S11a-c |
| FM3187 | FM3168; *ADH1:OsTIR1-9myc::URA3* | This study | S11b,c |
| FM3173 | FM588; *VPS24:AID*:9myc::KanMX* | This study | S11d |
| FM3175 | FM2807; VPS24:AID*:9myc::KanMX | This study | S11d |
| FM3190 | FM3173; *ura3-52::ADH1p:OsTIR1:9myc::URA3* | This study | S11a,b,d |
| FM3191 | FM3175; *ura3-52::ADH1p:OsTIR1:9myc::URA3* | This study | S11b,d |
| FM3238 | *MATa bar1Δ leu2-3,112 ura3-52 his3-Δ200 trp1-Δ63 ade2-1 lys2-801; NUP49:mCherry::NatNT2; HTA2:eCFP::TRP1; SEC61:eYFP::KanMX; Δsnf7::URA* | This study | S13 |

[1] Semicolon (";") separates genetic modifications accomplished sequentially through transformation. Intermediate strains are omitted.

### SUPPLEMENTARY MOVIE

**Movie S1. Z-series and 3D reconstruction of a nuclear septum in a late-M *Δmsc1* cell.** This movie is related to the cell shown in Figure S2a,b. The three first columns show a walk through the 34 z planes (Sec61-eCFP, Nup49-mCherry, and a merge of both), and the last two columns show rotations of the 3D reconstruction (X and Y axis, respectively). Unlike Figure S2, the whole z-series throughout the cell is included, so the cortical ER is clearly visualized in the Sec61-eCFP channel.
